# Distinct role of TGN-resident clathrin adaptors for Rab5 activation in the TGN-endosome trafficking pathway

**DOI:** 10.1101/2023.03.27.534325

**Authors:** Makoto Nagano, Kaito Aoshima, Hiroki Shimamura, Daria Elisabeth Siekhaus, Junko Y. Toshima, Jiro Toshima

## Abstract

Clathrin-mediated vesicle trafficking plays central roles in the post-Golgi transport pathways from the *trans*-Golgi network (TGN) to endosomes. In yeast, two clathrin adaptors – AP-1 complex and GGA proteins (GGAs) – are predicted to generate distinct transport vesicles at the TGN, and epsin-related Ent3p/Ent5p act as accessories for these adaptors. Recently, we showed that vesicle transport from the TGN, rather than from the plasma membrane, is crucial for Rab5-mediated endosome formation, and that Ent3p/5p are crucial for this process, whereas AP-1 and GGAs are dispensable. However, these observations were incompatible with previous studies showing that these adaptors are required for Ent3p/5p recruitment to the TGN, and thus the overall mechanism responsible for regulation of Rab5 activity remains ambiguous. Here we investigated the functional relationships between clathrin adaptors in post-Golgi-mediated Rab5 activation. We were able to show that AP-1 disruption in *ent3*Δ/*5*Δ mutant impairs Rab5-GEF Vps9p transport to the Rab5 compartment, and severely reduces Rab5 activity. Additionally, GGAs, Golgi-resident PI4 kinase Pik1p and Rab11 GTPases Ypt31p/32p were found to have partially overlapping functions for recruitment of AP-1 and Ent3p/5p to the TGN. These findings suggest a distinct role of clathrin adaptors for Rab5 activation in the TGN-endosome trafficking pathway.

## Introduction

Clathrin-mediated endocytosis is the process by which cells internalize various molecules, such as membrane proteins and extracellular molecules, through clathrin-coated vesicles that bud off from the plasma membrane (Kaksonen and Roux, 2018). Once endocytic vesicles are internalized into the cytosol, they are rapidly targeted to the early/sorting compartment that sorts endocytic cargos to the plasma membrane for recycling or to lysosomes/vacuoles for degradation (Grant and Donaldson, 2009; Mellman, 1996). The fundamental mechanism of endosome trafficking via the endocytic pathway is well conserved among eukaryotic cells, but the structural features of the early/sorting compartment differ (Huotari and Helenius, 2011; Scott et al., 2014). In plants, the *trans*-Golgi network (TGN) serves the role of the compartment, whereas in mammalian cells it functions as an independent organelle distinct from the TGN (Contento and Bassham, 2012; Scott et al., 2014; Tooze and Hollinshead, 1991). The endosomal system of the budding yeast *Saccharomyces cerevisiae* has also been studied extensively, and like animal cells, it has been shown to include at least two types of endosomal compartment: an early endosome-like compartment containing Rab5 (Vps21p) and a late endosome-like compartment containing Rab7 (Cabrera et al., 2014; Hutagalung and Novick, 2011; Shimamura et al., 2019; Toshima et al., 2014). On the other hand, a previous study has shown that yeast lacks distinct early endosomes, and that the TGN functions as an early endosome-like sorting compartment, as is the case in plant cell (Day et al., 2018). Additionally, our recent study revealed that the TGN sub-compartment (Tlg2p-residing compartment), in which a yeast syntaxin homologue Tlg2p resides, functions as the early/sorting compartment, sorting endocytic cargo to the Vps21p-residing compartments (Toshima et al., 2022). Thus, the identity of the Vps21p-residing compartment and how the compartment is generated is still under debate.

It has been suggested that Vps21p-residing endosomes, as well as early endosomes in animal cells, are formed and maintained by fusion of endocytic vesicles derived from the plasma membrane, and then mature into late endosomes, which receive TGN-derived vesicles (Huotari and Helenius, 2011; Scott et al., 2014; Shideler et al., 2015). However, in a recent study we demonstrated that although endocytic vesicle internalization from the plasma membrane is not essential for Vps21p-mediated endosome formation and trafficking, vesicle transport from the TGN plays a crucial role (Nagano et al., 2019). We also demonstrated that recruitment of Rab5 GEF Vps9p to the TGN is important for activation of Vps21p at the endosomal compartment (Nagano et al., 2019). A similar mechanism whereby Rab5 GEF is recruited to the TGN and activates Rab5 has been reported in plant cells (Minamino and Ueda, 2019), suggesting that the mechanism of Rab5 activation at the TGN is conserved between yeast and plant cells.

In yeast activation of Vps21p is partially suppressed by deletion of the epsin-like proteins, Ent3p/5p (Duncan et al., 2003a; Nagano et al., 2019). Ent3p/5p have been identified as accessory molecules that function with two major clathrin adaptors (Čopič et al., 2007; Costaguta et al., 2006; Daboussi et al., 2012): the AP-1 complex (AP-1) and GGAs (Gga1p/2p) (Traub, 2005). GGAs and AP-1 form two distinct clathrin-coated vesicles, the GGA-enriched vesicle (GGA vesicle), including Ent3p and a minor population of Ent5p, and the AP-1-enriched vesicle (AP-1 vesicle), which includes most of Ent5p and resides at the late TGN (Daboussi et al., 2012; Tojima et al., 2019). GGA vesicles are formed approximately 10 s prior to the AP-1 vesicles, and GGA vesicles mediate the transport of cargos from the TGN to the vacuole (Casler and Glick, 2020), whereas AP-1 vesicles mediate intra-Golgi recycling (Casler et al., 2021).

Curiously, in contrast to the deletion of Ent3p/5p, which affects Vps21p activity, deletion of GGAs or AP-1 complex, or even both, has a negligible effect on Vps21p activity (Nagano et al., 2019). These were seemingly contradictory observations because a series of previous studies focusing on the genetic and physical interactions between the clathrin adaptors indicated the functional relationships between the adaptors. For instance, GGAs are required for Ent3p recruitment to the TGN (Costaguta et al., 2006) and AP-1 has overlapping functions with Ent5p (Casler et al., 2021). These observations obscured the role of clathrin-mediated vesicle transport in Vps21p activation, and thus it is important to clarify which molecules, and the factor(s) that integrate them, are required for Vps21p activation in order to fully understand the mechanism regulating endosome formation and trafficking.

Several upstream factors that recruit these clathrin adaptors and regulate the progressive formation of GGA and AP-1 vesicles have been clarified. Phosphatidylinositol-4-phosphate (PI(4)P) (Daboussi et al., 2012; Wang et al., 2003), which is generated by Golgi-resident PI 4-kinase Pik1p (Walch-Solimena and Novick, 1999) is reported to play a central role in this process. Pik1p is initially recruited to the TGN through interaction with Arf1 GTPase and yeast Frequenin homologue Frq1p (Daboussi et al., 2012; Hendricks et al., 1999; Highland and Fromme, 2021). PI(4)P generated by Pik1p increases GGA localization at the TGN, and GGA in turn further recruits Pik1p (Daboussi et al., 2012; Daboussi et al., 2017; Highland and Fromme, 2021; Zhdankina et al., 2001). These positive feedback mechanisms facilitate the formation of GGA vesicles at an earlier TGN stage relative to the formation of AP-1 vesicles, and simultaneously induce localization of AP-1 and Ent5p at the TGN, leading to sequential formation of AP-1 vesicles at the later stage of the TGN (Daboussi et al., 2012). The yeast Rab11s, Ypt31p/32p, are other important regulators implicated in formation of transport carriers at the TGN (Benli et al., 1996; McDonold and Fromme, 2014; Ortiz et al., 2002). Previous studies have demonstrated a functional relationship between Arf1p and Ypt31p/32p in Pik1p-mediated PI(4)P production, and following vesicle formation at the TGN (Thomas and Fromme, 2016). Ypt31p/32p regulate the localization of Ent3p/5p at the TGN (Nagano et al., 2019). Thus, Pik1p and yeast Rab11s might be key regulators of Vps21p-mediated endosome formation.

In the present study, we aimed to clarify the functional relationships between clathrin adaptors in the Rab5 activation in yeast. To determine more accurately the level of Rab5 activity in the cell, we developed a novel biochemical method capable of detecting Rab5 activity with high sensitivity. Using this novel method and live-cell imaging, we were able to show that Ent3p/5p play a central role in the targeting of Vps9p from the TGN to endosomes, and that AP-1 cannot compensate this function. However, the AP-1 plays an important role for the retention of Vps9p at the TGN in the absence of Ent3p/5p. On the other hand, GGAs are involved in the Rab5 activation via the recruitment of Ent3p to the TGN. Together with GGAs, Pik1p and Ypt31p/32p have partially overlapping roles in the recruitment of Ent3p/5p and AP-1 to the TGN, and thereby these three factors act cooperatively in Vps21p activation. Thus, each of the three types of clathrin adaptors, Ent3p/5p, AP-1 and GGAs, contribute individually to Rab5 activation.

## Results

### AP-1 complex is involved in formation of the Vps21p-residing compartment

AP-1 and GGAs have been well-characterized as major clathrin adaptors, which regulate the clathrin-mediated vesicle transport from the TGN to endosome (Costaguta et al., 2001; Costaguta et al., 2006; Daboussi et al., 2012; Duncan et al., 2003a) and epsin-like Ent3p/5p have been identified as their accessory proteins (Costaguta et al., 2001; Costaguta et al., 2006; Daboussi et al., 2012; Duncan et al., 2003b). We have previously reported that Ent3p/5p is important for formation of the Vps21p-residing compartment (Vps21p compartment) (Nagano et al., 2019). Since previous studies have demonstrated that Gga1p/2p are involved in Ent3p, but not Ent5p, localization at the TGN, whereas AP-1 is dispensable for the localization of both (Costaguta et al., 2006), we first confirmed the role of these adaptor proteins in formation of the Vps21p compartment, using a synthetic genetic approach involving *ent3*Δ or *ent5*Δ mutation with deletion of the *GGA1*/*2* genes or *APL4* gene, encoding the AP-1γ subunit. To precisely evaluate the differences in the localization of Vps21p, each mutant was compared directly alongside the parental wild-type cells that had been labeled for their expression of Vph1-mCherry (Fig. 1A) or Hse1-tdTomato (Fig. 1B). In a previous study, we had demonstrated that the *ent3*Δ *ent5*Δ mutant shows decreased fluorescence intensity for GFP-Vps21p and an increase in the number of Vps21p compartments as a result of reduced Vps21p activity (Nagano et al., 2019). In both the *ent3*Δ *apl4*Δ and *ent5*Δ *apl4*Δ mutants, the number and fluorescence intensity of GFP-Vps21p-residing compartments were similar to those in wild-type cells (Fig. 1A, C, F), consistent with a previous observation that AP-1 is not necessary for Ent3p/5p localization (Costaguta et al., 2006). In the *gga1*Δ*/2*Δ mutant, there was a decrease of GFP-Vps21p fluorescence intensity (∼76%) and an increase in the number of Vps21p compartments when combined with the *ent5*Δ mutant (Fig. 1B, C, F), suggesting that GGA-dependent recruitment of Ent3p is important for formation of the Vps21p compartment.

**Figure 1.**
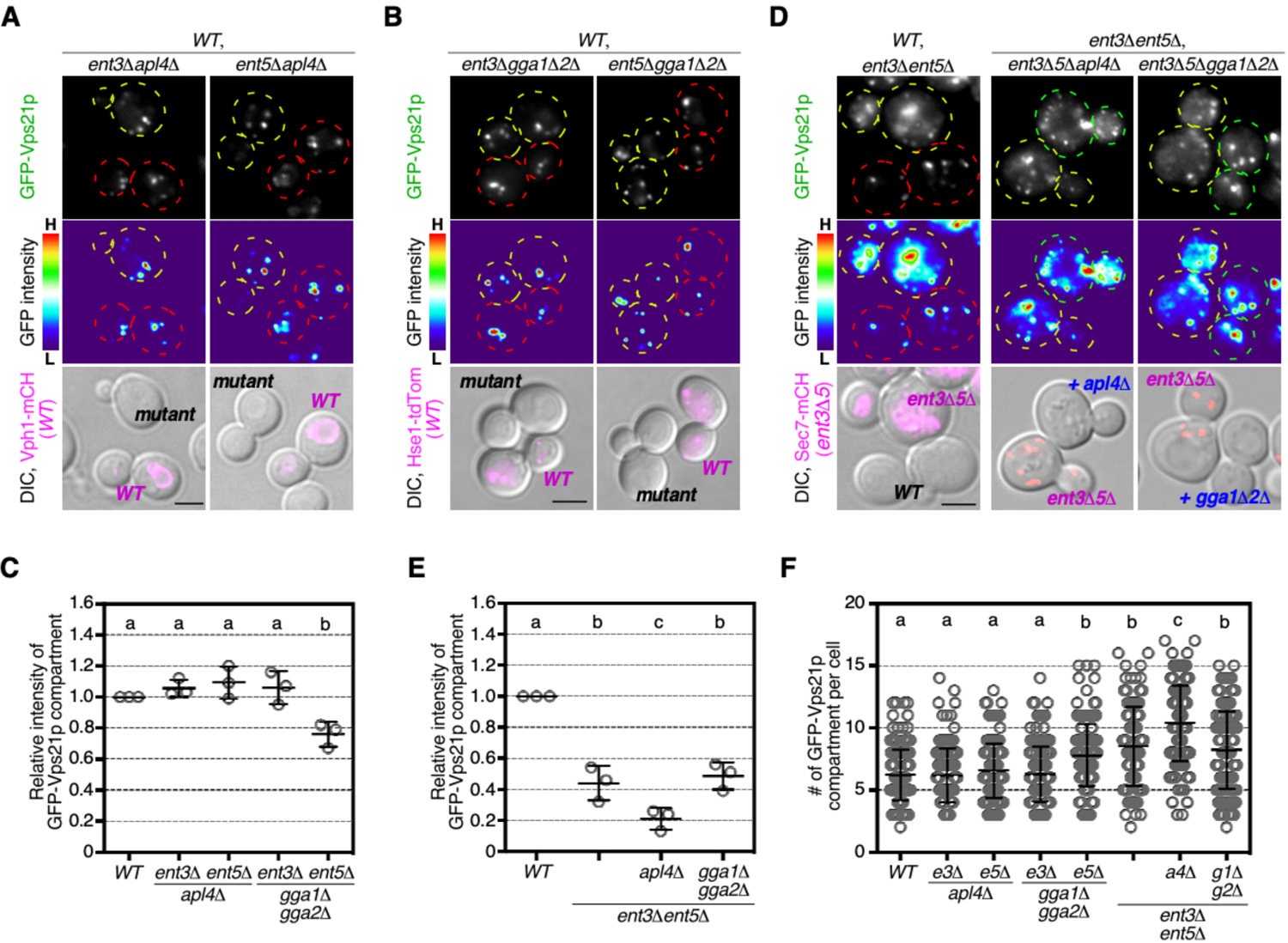
AP-1 complex disruption impairs the Vps21p-residing compartment formation in *ent3*Δ *ent5*Δ cells. (**A**) Localization of GFP-Vps21p in the *ent3*Δ *apl4*Δ or *ent5*Δ *apl4*Δ cells. The cells expressing GFP-Vps21p were grown to early-logarithmic to mid-logarithmic phase, mixed, and acquired in the same images. Fluorescence images or heat maps showing GFP levels are shown in the panels labeled GFP-Vps21p or GFP intensity, respectively. The wild-type (*WT*) cells are labeled by the expression of Vph1-mCherry (Vph1-mCH) which are shown in the lower images overlaid with DIC images. Red or yellow outline indicate the wild-type or mutant cells, respectively. (**B**) Localization of GFP-Vps21p in the *ent3*Δ *gga1*Δ *gga2*Δ or *ent5*Δ *gga1*Δ *gga2*Δ cells. Fluorescence images or heat maps showing GFP levels are shown in the panels labeled GFP-Vps21p or GFP intensity, respectively. The wild-type (*WT*) cells are labeled by the expression of Hse1-tdTomato (Hse1-tdTom) which are shown in the lower images overlaid with DIC images. Red or yellow outline indicate the wild-type or mutant cells, respectively. (**C**) Quantification of the fluorescence intensity of GFP-Vps21p-residing compartments displayed in (A) and (B). (**D**) Localization of GFP-Vps21p in the *ent3*Δ *ent5*Δ, *ent3*Δ *ent5*Δ *apl4*Δ, or *ent3*Δ *ent5*Δ *gga1*Δ *gga2*Δ cells. Fluorescence images or heat maps showing GFP levels are shown in the panels labeled GFP-Vps21p or GFP intensity, respectively. The wild-type (*WT*) cells in the left panels or *ent3*Δ *ent5*Δ mutant in the middle and right panels are labeled by the expression of Sec7-mCH which are shown in the lower images overlaid with DIC images. Red, yellow, or green outline indicate the *wild-type*, *ent3*Δ *ent5*Δ or other mutant cells, respectively. (**E**) Quantification of the fluorescence intensity of GFP-Vps21p-residing compartments displayed in (D). (**F**) Quantification of the number of GFP-Vps21p-residing compartments displayed in (A), (B) and (D). The number in the *wild-type* cells were quantified using the images displayed in (A). Data show mean ± SEM from three independent experiments in which 100 endosomes were scored per each experiment (C, E), or mean ± SD with 150 cells from three independent experiments (F). Different letters indicate significant difference at p < 0.05, one-way ANOVA with Tukey’s post-hoc test. Scale bar, 2.5 μm. *gga1*Δ*2*Δ: *gga1*Δ *gga2*Δ cells, *ent3*Δ*5*Δ: *ent3*Δ *ent5*Δ cells, *e3*Δ: *ent3*Δ cells, *e5*Δ: *ent5*Δ cells, *a4*Δ: *apl4*Δ cells, *g1*Δ*g2*Δ: *gga1*Δ *gga2*Δ cells.

Ent3p/5p is also required for Vps21p activity, and deletion of the *ENT3*/*5* genes decreases it to ∼36% of the level in wild-type cells (Nagano et al., 2019). This suggests that a factor other than Ent3p/5p is involved in Vps21p activation. To investigate whether the clathrin adaptors have additional roles in Vps21p compartment formation, we examined the localization of Vps21p in the *ent3*Δ *ent5*Δ mutant lacking *GGA1*/*2* or *APL4*. The *ent3*Δ *ent5*Δ *gga1*Δ*/2*Δ mutant exhibited Vps21p localization similar to that in the *ent3*Δ *ent5*Δ mutant, indicating that combination of *ent3/5*Δ and *gga1/2*Δ does not have an additive effect on Vps21p localization (Fig. 1D, E, F). Interestingly, we found that in the *ent3*Δ *ent5*Δ *apl4*Δ mutant the fluorescence intensity of the GFP-Vps21p compartments was further decreased relative to that in *ent3*Δ *ent5*Δ mutants, and that the number of GFP-Vps21p-labeled endosomes was also significantly increased (Fig. 1D-F). These results suggest that AP-1, as well as Ent3p/5p, has a role in formation of the Vps21p compartment.

### Deletion of AP-1 complex increases the inactivation state of Vps21p in the *ent3*Δ *ent5*Δ mutant

In a previous study employing a pull-down assay using the Rab5 effector, we showed that the activity of Vps21p in *ent3*Δ *ent5*Δ mutants was significantly decreased in comparison with that in wild-type cells (Nagano et al., 2019). Since the *ent3*Δ *ent5*Δ *apl4*Δ mutant exhibited severer defect in formation of the Vps21p compartment, we speculated that the activity of Vps21p is further decreased in the mutant. To assess with high precision what % of Vps21p activity is decreased in the mutant, we developed a new method with improved sensitivity and quantitative accuracy to assess the levels of active Rab5 (Fig. 2A). We utilized Nano luciferase (Boute et al., 2016) fused to either a nanobody that recognizes GFP (Katoh et al., 2015; Katoh et al., 2016) (Nluc-GNB) to detect total protein or to a fragment of human Rabenosyn-5 (Qi et al., 2015) that binds specifically to the GTP-bound form of Vps21p (Nluc-RbNT) (Fig. S1A-C). We confirmed that Nluc-GNB and Nluc-RbNT had almost the same enzyme activity (Fig. S1D). We tagged Vps21p with ALFA peptide (Götzke et al., 2019) and confirmed that GFP-ALFA-Vps21p showed a similar intracellular localization to GFP-Vps21p in wild-type cells and the *vps9*Δ *muk1*Δ mutant, which lacks Vps21p GEFs (Paulsel et al., 2013) (Fig. S1E). We also tested to be sure that GFP-ALFA-Vps21p, but not GFP-Vps21p, was specifically pulled down from the cell extracts using the GST-fused ALFA nanobody (Götzke et al., 2019) (Fig. S1F, G). GFP-ALFA-Vps21p was expressed in both strains at a similar level, although two bands with different molecular weights were detected only in the *vps9*Δ *muk1*Δ mutant (Fig. S1G). Consistent with this, when the pulled down GFP-ALFA-Vps21p was incubated with Nluc-GNB the luminescence signal reflecting the total amount of Vps21p was almost the same in wild-type and *vps9*Δ *muk1*Δ cells, whereas that of Nluc-RbNT, reflecting the amount of active Vps21p, was only detected in the pulled-down fraction from wild-type cells and not in that from the mutant (Fig. 2B). We normalized the Nluc-RbNT signals with the Nluc-GNB signals to determine the amount of active Vps21p (Fig. 2C). This result indicated that the majority of Vps21p exists as an inactive form bound to GDP in the *vps9*Δ *muk1*Δ mutant, consistent with the observation that Vps21p rarely localized at endosomal compartments in the mutant (Fig. S1E). With this newly developed method, we next examined the levels of activated Vps21p in the *ent3*Δ *ent5*Δ or *ent3*Δ *ent5*Δ *apl4*Δ mutant. We confirmed that GFP-ALFA-Vps21p was expressed in both strains at a similar level (Fig. S1H). The luminescence signals of Nluc-RbNT were significantly decreased in both *ent3*Δ *ent5*Δ and *ent3*Δ *ent5*Δ *apl4*Δ mutants, compared to wild-type cells, while the total amounts of Vps21p were almost the same (Fig. 2D). The level of activated Vps21p normalized by the RbNT/GNB ratio decreased to ∼45% or ∼10% in *ent3*Δ *ent5*Δ or *ent3*Δ *ent5*Δ *apl4*Δ mutants, compared to that in wild-type cells (Fig. 2E), indicating that deletion of AP-1 increased the inactivation state of Vps21p in the *ent3*Δ *ent5*Δ mutant. These results suggest that AP-1 functions redundantly with Ent3p/5p in Vps21p activation.

**Figure 2.**
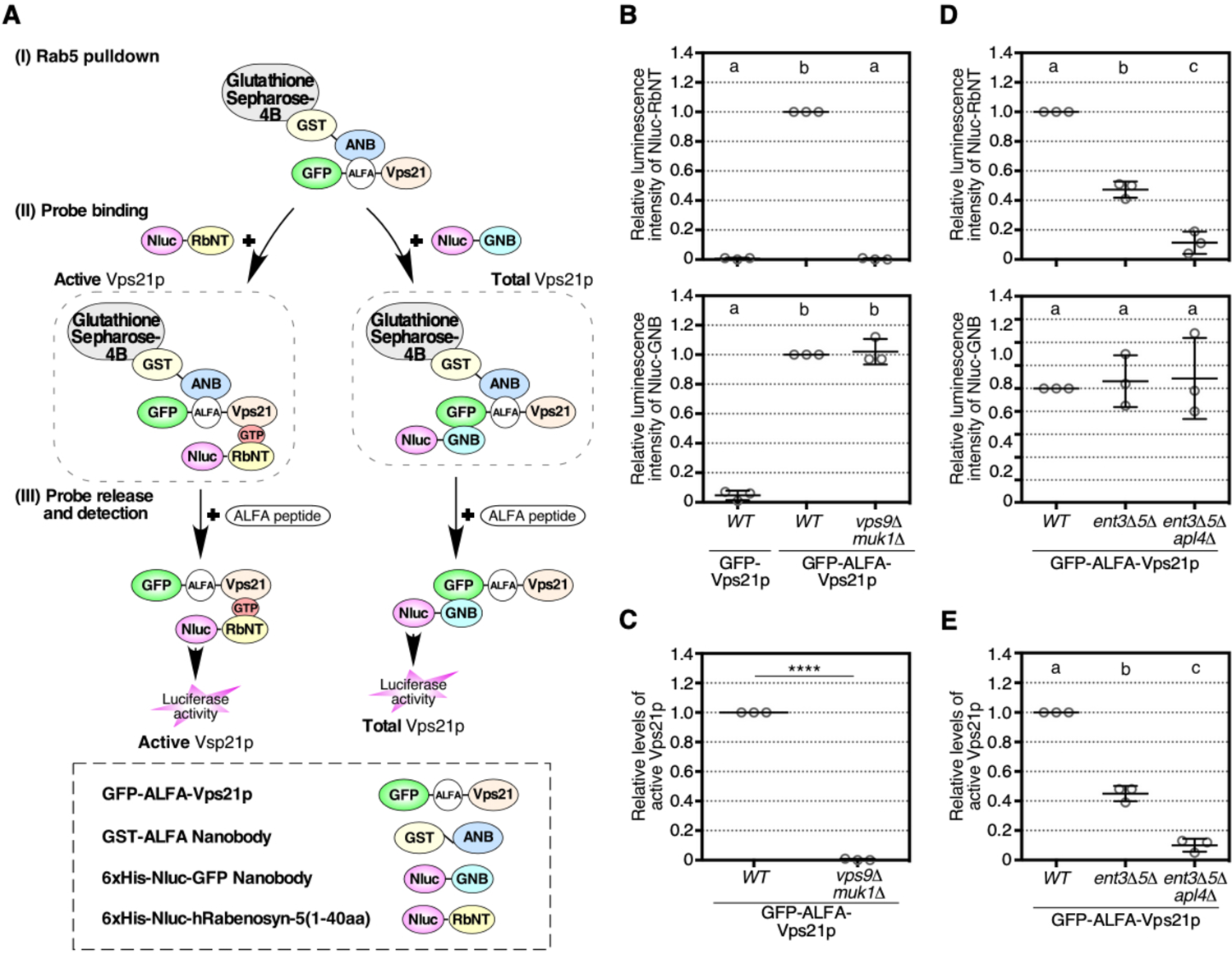
AP-1 complex disruption decreases the active state of Vps21p in *ent3*Δ *ent5*Δ cells. (**A**) Schematic illustration of the quantification assay for the intracellular active Rab5 levels. GFP-ALFA-Vps21p was purified from the yeast cell lysate using GST-ALFA nanobody (GST-ANB)-bound Glutathione sepharose-4B (i). The sepharose was equally divided into two fractions, and then added 6×His-Nano luciferase (Nluc)-Rabenosyn5 N-terminal fragment (Nluc-RbNT), a probe binding to the active Vps21p specifically, or 6×His-Nluc-GFP nanobody (Nluc-GNB), a probe binding to GFP (ii). After washout unbound probes, the bound probes were eluted by ALFA peptide, and then quantify the Nluc activity in the fraction (iii) (See in Materials and Methods). (**B, D**) Quantification of the binding amount of Nluc-fused probes in the total or active Vps21p pulldown fraction. GFP-Vps21p or GFP-ALFA-Vps21p was expressed in the indicated cells. According to the protocol illustrated in (A), Nluc activity in the Nluc-RbNT-bound active Vps21p fraction (upper graph) or the Nluc-GNB-bound total Vps21p fraction (lower graph) were quantified. (**C, E**) Quantification of the active Vps21p levels in the cells. Ratio of Nluc activity from Nluc-RbNT to that from Nluc-GNB in (B) or (D) was calculated as the active Vps21p levels in the cells. Graphs represent the active Vps21p level in the mutant cells relative to that in *WT* cells. Data show mean ± SEM from three independent experiments. Different letters indicate significant difference at p < 0.05, one-way ANOVA with Tukey’s post-hoc test (B, D, E). ****p < 0.0001, unpaired t-test with Welch’s correction (C).

### Endosomal fusion is impaired in the *ent3*Δ *ent5*Δ *apl4*Δ mutants

Previous studies of a mutant lacking *VPS9*, encoding a major Rab5 GEF, revealed accumulation of 40- to 50-nm vesicles or vesicle clusters proximal to the vacuole that were unable to fuse with each other (Burd et al., 1996; Cabrera et al., 2013). Other studies also demonstrated accumulation of Vps21p proximal to the vacuole in the *vps9*Δ mutant (Markgraf et al., 2009). In accord with these observations, we found accumulation of the GFP-Vps21p signal in the *vps9*Δ mutant (Fig. 3A, B). This aberrant GFP-Vps21p signal was also observed in the *ent3*Δ *ent5*Δ and *ent3*Δ *ent5*Δ *apl4*Δ mutants, but not in wild-type cells (Fig. 3A, B). The accumulation of Vps21p appeared to correlate with the rate of Vps21p inactivation, because ∼80% of *vps9*Δ, ∼58% of *ent3*Δ *ent5*Δ *apl4*Δ and ∼15% of *ent3*Δ *ent5*Δ mutants exhibited this Vps21p accumulation (Fig. 3B). Next, using electron microscopy, we investigated whether the aberrant GFP-Vps21p signal was attributable to accumulation of vesicle structures. Similarly to *vps21*Δ *ypt52*Δ mutants, which lack the major yeast Rab5s, vesicles of various sizes (∼40-150 nm) were found to accumulate in *ent3*Δ *ent5*Δ *apl4*Δ mutants, whereas such structures were rarely observed in wild-type cells (Fig. 3C, D). We also found that ∼80-150-nm vesicles were clustered together adjacent to the vacuole (Fig. 3D). Similar vesicle structures were also observed in *vps9*Δ mutant (Burd et al., 1996), suggesting that endosomal fusion mediated by Vps21p is impaired in *ent3*Δ *ent5*Δ *apl4*Δ mutants.

**Figure 3.**
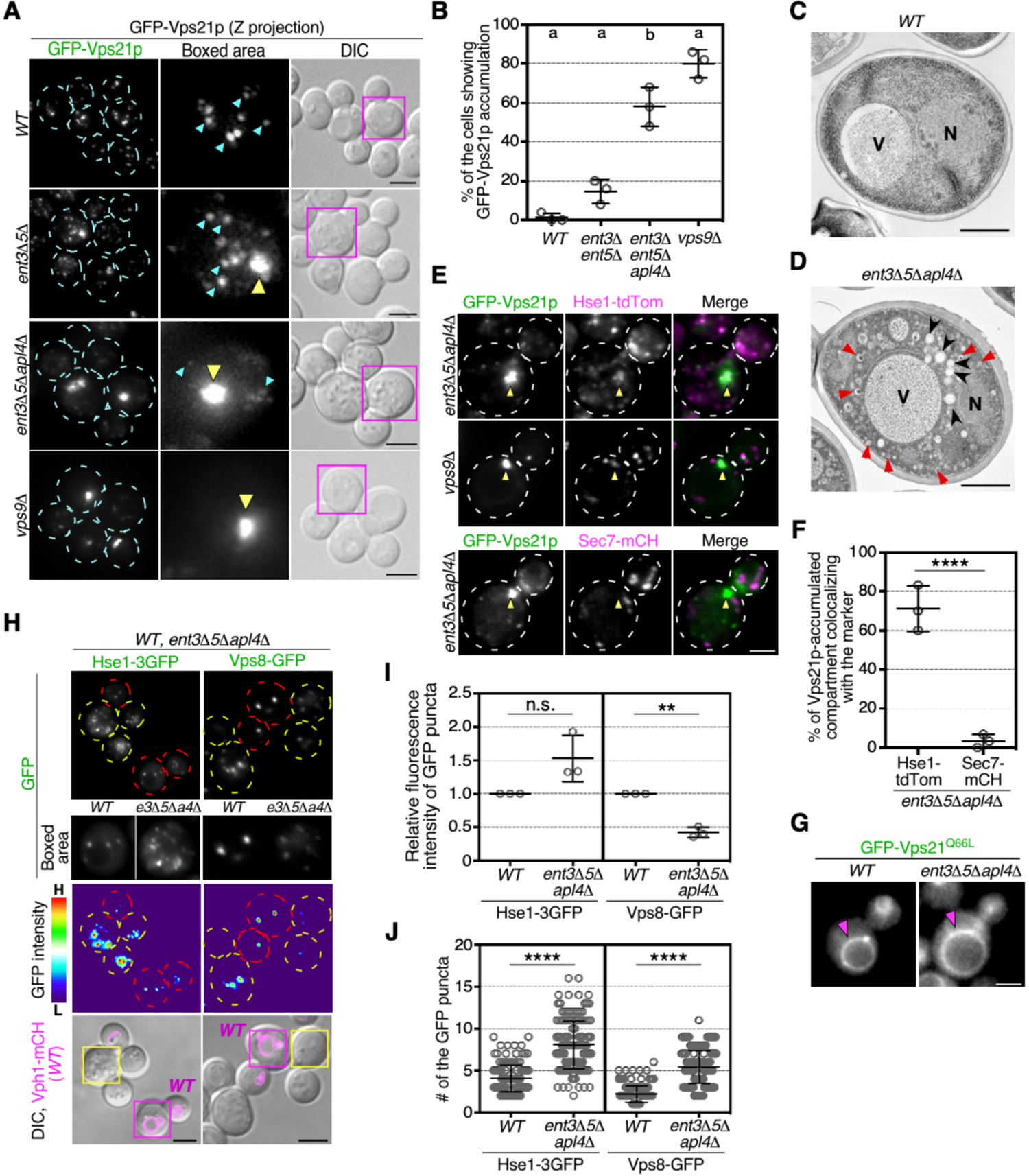
AP-1 complex disruption impairs the endosomal fusion in *ent3*Δ *ent5*Δ cells. (**A**) Maximum intensity projections of Z stacks of wild-type (*WT*) and mutant cells expressing GFP-Vps21p. The Z series was acquired through the entire cell at 0.4 μm intervals. Cyan and yellow arrowheads indicate the examples of GFP-Vps21p-residing endosome-like compartments and the GFP-Vps21p-accumulated regions, respectively. Higher magnification view of the boxed area is displayed in the middle panel. **(B)** The percentages of the cells showing the GFP-Vps21p accumulation displayed in (A). **(C, D)** Ultrastructure of the intracellular compartments observed in *wild-type* (C) and *ent3*Δ*5*Δ *apl4*Δ (D) cells. Cells were grown at 25 °C, fixed using propane jet freezing method and processed for electron microscopic analysis. Red arrowheads indicate examples of ∼50-nm vesicles, and black arrowheads indicate vesicle clusters proximal to the vacuole. N: nucleus, V: vacuole. (**E**) Co-localization of GFP-Vps21p and Hse1-tdTomato (tdTom) or Sec7-mCherry (mCH) in the *ent3*Δ*5*Δ *apl4*Δ or *vps9*Δ cells. Arrowheads indicate the example of GFP-Vps21p-accumulated compartments. **(F)** The percentages of the overlapping localization of GFP-Vps21p-accumulated compartments with the mCH/tdTom-tagged marker, as displayed in (E). **(G)** Localization of GFP-tagged constitutively active Vps21Q66L mutant in the cells. Arrowheads indicate the localization at the vacuolar membrane. (**H**) Localization of Hse1p tagged with three tandem repeats of GFP (Hse1-3GFP) or Vps8p tagged with GFP (Vps8-GFP) in the *ent3*Δ*5*Δ *apl4*Δ cells. The cells expressing the GFP-tagged marker were grown to early-logarithmic to mid-logarithmic phase, mixed, and acquired in the same images (left panels: Hse1-3GFP, right panels: Vps8-GFP). Fluorescence images or heat maps showing GFP levels are shown in the panels labeled GFP or GFP intensity, respectively. The wild-type cells are labeled by the expression of Vph1-mCherry (Vph1-mCH) which are shown in the lower images overlaid with DIC images. The boxed areas are shown at higher magnification in the lower side in the GFP panels. Red or yellow outline indicate the wild-type or mutant cells, respectively. (**I**) Quantification of the fluorescence intensity of Hse1-3GFP (left graph) or Vps8-GFP (right graph) displayed in (H). (**J**) Quantification of the number of Hse1-3GFP (left graph) or Vps8-GFP (right graph) puncta displayed in (H). Data show mean ± SEM from three independent experiments in which 50 cells (B), 30 compartment (F) or 50 puncta (I) were scored per each experiment, or mean ± SD with 150 cells from three independent experiments (J). Different letters indicate significant difference at p < 0.05, one-way ANOVA with Tukey’s post-hoc test (B). n.s., not significant, **p < 0.01, ****p < 0.0001, unpaired *t*-test with Welch’s correction (F, I, J). Scale bars: 2.5 μm (A, E, H, G), 1 μm (C, D). *e3*Δ*5*Δ*a4*Δ: *ent3*Δ *ent5*Δ *apl4*Δ cells.

By comparing the localization of Vps21p with mCherry- or tdTomato-tagged specific markers for organelles, we found that Hse1p, a marker of the endosome (Bilodeau et al., 2002; Toshima et al., 2014), highly localizes (∼71.1%), but Sec7p, a marker of the TGN (Achstetter et al., 1988), rarely localizes (∼3.3%) at the Vps21p cluster adjacently to the vacuole (Fig. 3E, F), suggesting that the vesicles accumulating adjacently to the vacuole in the *ent3*Δ *ent5*Δ *apl4*Δ mutant has endosomal characteristics. This aberrant GFP-Vps21p accumulation in the *ent3*Δ *ent5*Δ *apl4*Δ mutant was suppressed by expression of the constitutively active Q66L mutant of Vps21p (Fig. 3G). These observations suggested that the vesicle clusters in *ent3*Δ *ent5*Δ *apl4*Δ were formed by impairment of the endosome fusion steps regulated by Rab5. We next sought to determine whether *ent3*Δ *ent5*Δ *apl4*Δ affects trafficking processes across the entire mechanism of endosome formation/maturation or at specific step(s) including Rab5 activation. To this end, we visualized two of the endosome-resident proteins, Hse1p and Vps8p, as a GFP-tagged form in wild-type and *ent3*Δ *ent5*Δ *apl4*Δ cells. Hse1p, an ESCRT-0 subunit, resides at the endosomal compartment via the ubiquitin-binding domain (Bilodeau et al., 2002), whereas Vps8p localizes at the PVC in a Rab5-dependent manner (Cabrera et al., 2013). Interestingly, the fluorescence intensity of GFP-tagged Hse1p at the punctate compartment was slightly increased in the *ent3*Δ *ent5*Δ *apl4*Δ mutant, even though that of GFP-tagged Vps8p was decreased, similarly to GFP-Vps21p (Fig. 3H, I). Additionally, Hse1-3GFP and Vps8-GFP puncta were significantly increased in the *ent3*Δ *ent5*Δ *apl4*Δ mutant (Fig. 3H, J). These observations indicated that Ent3p/5p and AP-1 are required for formation of the Vps21p compartment, followed by fusion between the early endosomal compartments regulated by Rab5 effectors including CORVET, but dispensable for recruitment of the ESCRT-0 subunit to the compartments.

### GGAs act redundantly with Pik1p and Ypt31p/32p upstream of Ent3p, thereby contributing to Vps21p activation

We then sought to identify upstream factors involved in the recruitment of Ent3p/5p and AP-1 to the TGN in Rab5 activation. Although Gga1p/2p has been shown to be crucial for localization of Ent3p at the TGN (Costaguta et al., 2006), the effect on Vps21p localization in the *gga1*Δ/*2*Δ *ent5*Δ mutant (∼76%) (Fig. 1C) was less severe than that in *ent3*Δ *ent5*Δ (∼44%) (Fig. 1E), suggesting the involvement of other factors in Ent3p recruitment. Two possible factors that might regulate the localization of Ent3p/5p and AP-1 at the TGN are PI(4)P and yeast Rab11 homologues Ypt31p/32p. Previous studies have demonstrated that PI(4)P generated by TGN-resident PI 4-kinase Pik1p is crucial for recruitment of Ent5p and AP-1 (Daboussi et al., 2012; Daboussi et al., 2017; Highland and Fromme, 2021), and that Ypt31p/32p is also involved in localization of Ent3p/5p to the TGN (Nagano et al., 2019). Therefore, we examined the contribution of GGAs, Pik1p and Ypt31p/32p to recruitment of Ent3p/5p and AP-1 to the TGN. We first confirmed that deletion of the *GGA1* and *GGA2* genes significantly decreased the localization of Ent3-GFP at the TGN to ∼33% of wild-type cell, without changing the localization of Ent5-GFP and Apl2-GFP (Fig. 4A, and Fig. S2A, B). Next, to investigate the contribution of PI(4)P, we used a temperature-sensitive *pik1-1*(D1055G) mutant, in which the PI(4)P level at the TGN was decreased to ∼6% of wild-type cells at the non-permissive temperature (37°C) (Yamamoto et al., 2017). The fluorescence intensity of Sec7-mCherry at the TGN was not affected in the *pik1-1* mutant (Fig. S2C, D). In the *pik1-1* mutant, in agreement with previous reports (Daboussi et al., 2012; Daboussi et al., 2017; Highland and Fromme, 2021), the fluorescence intensity of GFP-tagged Ent3p, Ent5p or Apl2p, which is a subunit of AP-1, at the TGN decreased to ∼53%, ∼21% or ∼23% of that in wild-type cells, respectively (Fig. 4A, and S2C, D). Our previous work on yeast Rab11 (Ypt31p/32p) had shown that in the *ypt31ts* mutant Ent3p and Ent5p were obviously less prevalent at the TGN at 37°C (∼33% and ∼28%, respectively), whereas AP-1 localization at the TGN was only partially reduced (∼83%) (Fig. 4A) (Nagano et al., 2019).

**Figure 4.**
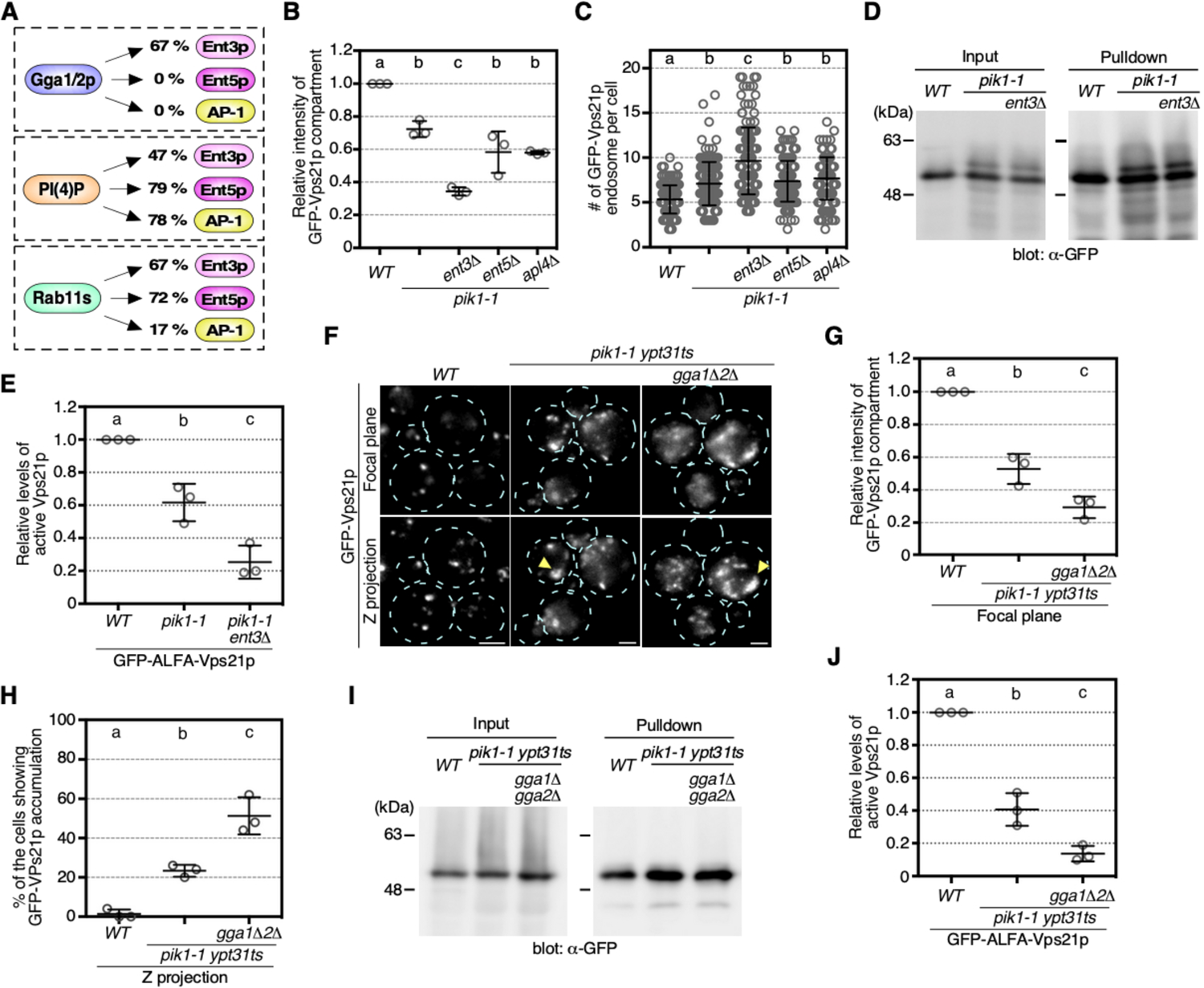
Gga1p/2p, Pik1p and Ypt31p/32p redundantly regulate Vps21p activity. (**A**) Summary of the contribution of Gga1p/2p, phosphatidylinositol 4-phosphate generated by Pik1p, or Rab11 homologues Ypt31p/32p to the TGN localization of Ent3p/5p and AP-1 complex. **(B, C)** Quantification of the fluorescence intensity (B) or number (C) of GFP-Vps21p-residing compartments displayed in Fig. S3A. **(D)** Immunoblots showing the amount of GFP-ALFA-Vps21p in the cell extract (Input) or the pulldown (Pulldown) fraction in Fig. S3D. (**E**) Quantification of the active Vps21p levels in the cells. The ratio of Nluc activity from Nluc-RbNT to that from Nluc-GNB in Fig. S3D was defined as the active Vps21p levels in the cells. Graphs represent the active Vps21p level in the mutant cells relative to that in *WT* cells. (**F**) Focal plane images and maximum intensity projections of Z stacks of the *WT* and mutant cells expressing GFP-Vps21p. The Z series was acquired through the entire cell at 0.4 μm intervals. Yellow arrowheads indicate the example of the GFP-Vps21p-accumulated regions. (**G**) Quantification of the fluorescence intensity of GFP-Vps21p puncta in the focal plane images displayed in (F). (**H**) The percentages of the cells showing GFP-Vps21p accumulation in the projections of Z stacks displayed in (F). **(I)** Immunoblots showing the amount of GFP-ALFA-Vps21p in the cell extract (Input) or the pulldown (Pulldown) fraction in Fig. S3E. (**J**) Quantification of the active Vps21p levels in the cells. The ratio of Nluc activity from Nluc-RbNT to that from Nluc-GNB in Fig. S3E was defined as the active Vps21p levels in the cells. Graphs represent the active Vps21p level in the mutant cells relative to that in *WT* cells. Data show mean ± SEM from three independent experiments in which 100 compartments (B), 100 puncta (G) or 50 cells (H) were scored per each experiment, mean ± SEM from three independent experiments (E, J), or mean ± SD with 150 cells from three independent experiments (C). Data show Different letters indicate significant difference at p < 0.05, one-way ANOVA with Tukey’s post-hoc test. Scale bars: 2.5 μm

We next examined the effect of reduction of these upstream factors on Vps21p compartments. By directly comparing the *pik1-1* mutant with wild-type cells, we found that the fluorescence intensity of GFP-Vps21p-residing endosomes in the mutant decreased to ∼72% of that in wild-type cells (Fig. 4B and Fig. S3A), and that the number of endosomes was increased relative to wild-type cells (Fig. 4C and Fig. S3A), indicating that a decreased PI(4)P level reduces Vps21p activity and thus inhibits fusion of Vps21p compartments. Deletion of the *ENT3* gene in the *pik1-1* mutant further decreased Vps21p localization at the endosome and increased the number of endosomes, whereas deletion of *ENT5* or *APL4* in the *pik1-1* mutant had a negligible effect on them (Fig. 4B, C, and Fig. S3A), consistent with the PI(4)P’s contribution to the recruitment of Ent5p and AP-1 (Daboussi et al., 2012; Daboussi et al., 2017; Highland and Fromme, 2021). Additionally, in the *pik1-1 ent3*Δ mutants, the GFP-Vps21p accumulation adjacent to the vacuole was observed in the same proportion of cells as in the *ent3*Δ *ent5*Δ *apl4*Δ mutant, but was only slightly present in other mutants (Fig. S3B, C). Furthermore, we found that the GTP-bound active form of Vps21p was significantly decreased to 62% in *pik1-1* mutants, and further decreased to 22% in the *pik1-1 ent3*Δ mutants, relative to wild-type cells (Fig. 4D, E, and Fig. S3D). These results suggest that PI(4)P is a key player for the Vps21p activation, but PI(4)P-independent recruitment of Ent3p to the TGN is also important for the process.

Since each of PI(4)P, Rab11, and GGAs are involved in the recruitment of Ent3p to the TGN, we next examined whether they function in the Rab5 activation in parallel or in cooperation. The fluorescence intensity of GFP-Vps21p at the puncta in the *pik1-1 ypt31ts* mutant (∼53%) was lower than that in the *pik1-1* mutant, and even lower in the *pik1-1 ypt31ts gga1*Δ*2*Δ mutant (∼29%) (Fig. 4F, G). The GFP-Vps21p accumulation was observed in ∼23% of the *pik1-1 ypt31ts* mutant and ∼51% of the *pik1-1 ypt31ts gga1*Δ*2*Δ mutant (Fig. 4H). Consistent with these observations, the activity of Vps21p in the *pik1-1 ypt31ts gga1*Δ *gga2*Δ mutant (14%) was further decreased, relative to that in the *pik1-1 ypt31ts* mutant (41%) (Fig. 4I, J, and Fig. S3E), suggesting that these upstream factors cooperatively recruit Ent3p and contribute to the activation of Vps21p.

### AP-1 plays a role in maintaining Vps9p in the TGN–endosome trafficking pathway in the *ent3*Δ*5*Δ mutant

We next wished to examine the effect of deletion of AP-1 on Vps9p localization, since we have shown previously that transport of Vps9p from the TGN to the endosome is necessary for Vps21p activation (Nagano et al., 2019). Vps9p is localized at several puncta, which were identified as endosomes in the previous study, in addition to the cytosol in wild-type cells (Fig. 5A) (Nagano et al., 2019). In the *ent3*Δ *ent5*Δ mutant the number of GFP-Vps9p puncta was increased relative to that in wild-type cells (Fig. 5A, B), and the majority of Vps9p showed a change in localization from endosomes to the TGN (Nagano et al., 2019). Thus, we speculated that the punctuate localization of GFP-Vps9p would be further increased in the *ent3*Δ *ent5*Δ *apl4*Δ mutant. As expected, by directly comparing the mutants, we found an increased number of Vps9p puncta in the *ent3*Δ *ent5*Δ *apl4*Δ mutant relative to the *ent3*Δ *ent5*Δ mutant (Fig. 5A, B). To make Vps9p localization at endosomes clearer, we used a class E *vps* mutant, which contains a large aberrant endosomal structure called class E compartment (Paulsel et al., 2013). We expressed an mCherry-C-terminally tagged Snf7p, a subunit of ESCRT-III, which has been shown as a GFP fusion to exhibit a dominant-negative phenotype in wild-type cells, causing the formation of the Class E compartment (Teis et al., 2008). We observed Vps9p localizing to the enlarged compartment observed in wild-type cells expressing mCherry-fused Snf7p, (Fig. 5C, D). Interestingly, formation of the class E compartment in the *ent3*Δ *ent5*Δ mutant was decreased to ∼34% of that in wild-type cells, and was diminished almost completely in the *ent3*Δ *ent5*Δ *apl4*Δ mutant (Fig. 5C, E). Since activity of Vps21p is known to be required for formation of the class E compartment (Russell et al., 2012), these observations are consistent with our findings that most Vps21p exists in an inactive state in the *ent3*Δ *ent5*Δ *apl4*Δ mutant. Quantitative analysis revealed that GFP-Vps9p localization at class E compartments was decreased significantly to ∼29% in the *ent3*Δ *ent5*Δ mutant, and to 0% in the *ent3*Δ *ent5*Δ *apl4*Δ mutant (Fig. 5C, D). In the *ent3*Δ *ent5*Δ *apl4*Δ mutants, GFP-Vps9p was localized at small puncta, different from the late endosomal compartments where Snf7p was localized (Fig. 5C). These results suggest that in the *ent3*Δ *ent5*Δ *apl4*Δ mutant Vps9p localization at the Golgi might be increased whereas its localization at the endosome is completely lost. To determine the compartment to which GFP-Vps9p localizes in the *ent3*Δ *ent5*Δ *apl4*Δ mutants, we first compared its localization with Vps21p. To clearly detect the localization of Vps9p at small puncta, we reduced the cytoplasmic fluorescence background of Vps9p by the top-hat (Legland et al., 2016) and bilateral filters (Chaudhury et al., 2011) bundled in the Image J FIJI software package (Schindelin et al., 2012) (Fig. S4A). By comparing the raw data with the processed data, we confirmed that the punctate localization of Vps9p can be extracted accurately (Fig. S4B). Using these processed images, we examined the localization of GFP-Vps9p with mCherry-Vps21p in wild-type or mutant cells. In wild-type cells, Vps9p puncta were highly co-localized with Vps21p (∼74%), however the co-localization decreased to ∼49% or ∼27% in the *ent3*Δ *ent5*Δ or *ent3*Δ *ent5*Δ *apl4*Δ mutant, respectively (Fig. 5F, G, and Fig. S4C), consistent with the reduced levels of Vps21p activation we had observed in these genotypes (Fig. 2E). Thus, Vps21p seemed not to be recruited to the small puncta of Vps9p increased in the *ent3*Δ *ent5*Δ *apl4*Δ mutants.

**Figure 5.**
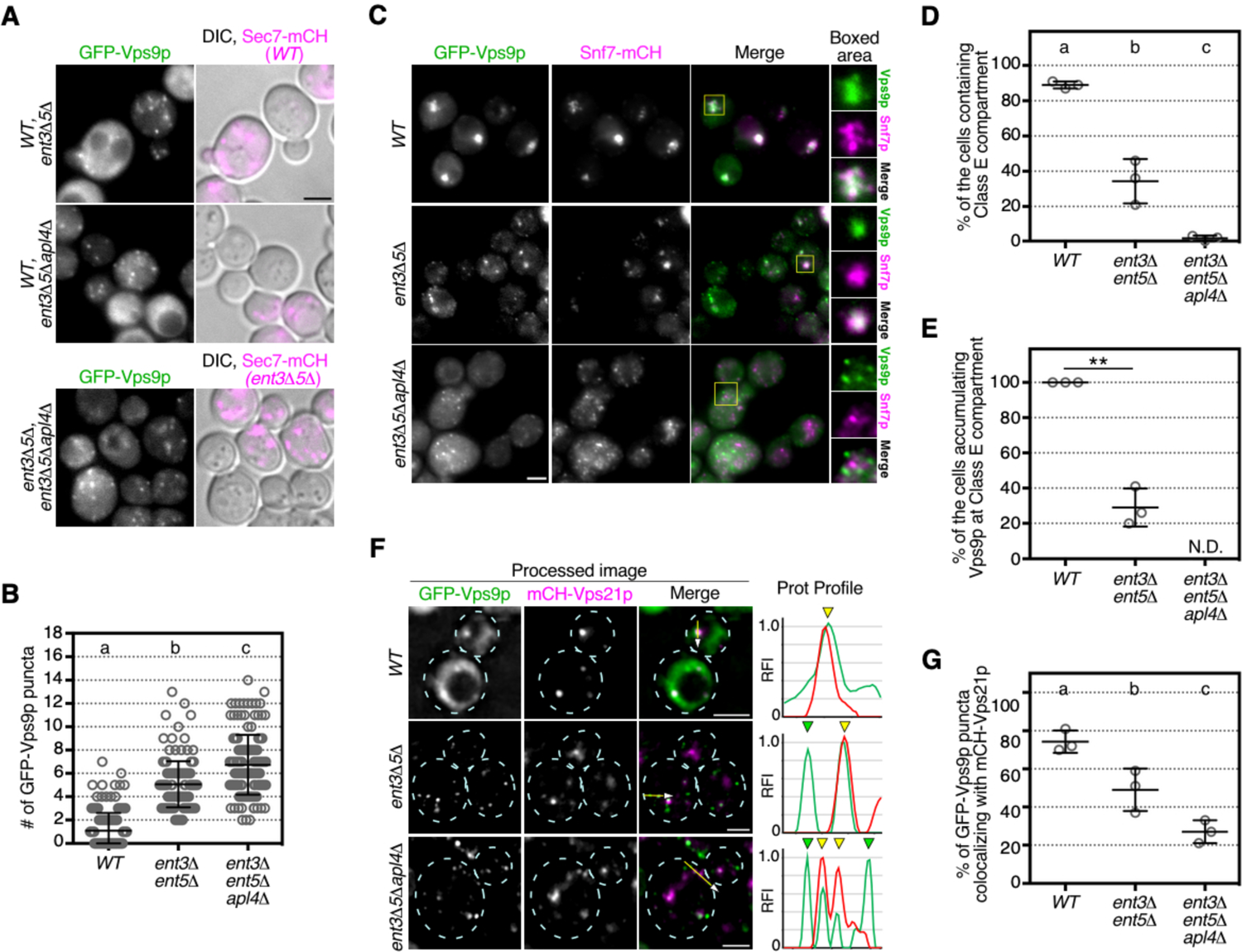
AP-1 complex disruption perturbs the targeting of Vps9p to endosomes in *ent3*Δ *ent5*Δ cells. (**A**) Localization of GFP-Vps9p in the *WT* and mutant cells. GFP-Vps9p was expressed under the control of the *ZWF1* promoter from the endogenous locus. The cells were grown to early-logarithmic to mid-logarithmic phase, mixed, and acquired in the same images. Fluorescence images of GFP-Vps9p are shown in the left panels. *WT* (upper and middle panels) or *ent3*Δ*5*Δ (lower panels) are labeled by the expression of Sec7-mCherry, which is shown in the images overlaid with DIC images in the lower panels. (**B**) Quantification of the number of GFP-Vps9p puncta displayed in (A). (**C**) Localization of GFP-Vps9p in the cells expressing Snf7-mCH. Snf7-mCH was expressed under the control of its own promoter in the endogenous locus. The boxed areas are shown at higher magnification in the right panels. (**D**) The percentages of the cells containing Class E compartments formed by Snf7-mCH accumulation displayed in (C). (**E**) The percentages of the cells accumulating GFP-Vps9p at Class E compartments formed by Snf7-mCH accumulation displayed in (C). (**F**) Dual color imaging of GFP-Vps9p and mCH-Vps21p in the *WT* and mutant cells. Fluorescence images of GFP-Vps9p and mCH-Vps21p shown in Fig. S4C were processed to reduce the cytoplasmic fluorescence background by sequential filtering using Top-hat and bilateral filters bundled in the Image J Fiji software package as shown in Fig. S4B. Representative fluorescence intensity profiles along an arrow are indicated in the right graphs. Yellow or green arrowheads indicate the presence or absence of the overlapping puncta, respectively. (**G**) Quantification of GFP-Vps9p puncta overlapping with mCH-Vps21p in the cells displayed in (F). Data show mean ± SD with 150 puncta from three independent experiments (B) or mean ± SEM from three independent experiments in which 50 cells (D, E) or 100 puncta (G) were scored per each experiment. Different letters indicate significant difference at p < 0.05, one-way ANOVA with Tukey’s post-hoc test (B, D, G). **p < 0.01, unpaired t-test with Welch’s correction (E). Scale bar, 2.5 μm. N.D., not determined. Scale bars: 2.5 μm

Based on this, we further attempted to reveal the compartment identity of the Vps9p puncta. By comparing the localization of GFP-Vps9p with specific markers for the TGN or endosomal compartments, we examined whether the dysfunction of AP-1 in the *ent3*Δ *ent5*Δ mutant further perturbs the targeting of Vps9p to endosomes. To this end, we employed Sec7-mCherry as a marker of the late TGN (Tojima et al., 2019), Gga2-tdTomato as a marker of the early TGN (Tlg2p-residing compartment) (Tojima et al., 2019; Toshima et al., 2022), Vps8-mCH as a marker of the Vps21p compartment (Fig. 6A). In wild-type cells, ∼8.7% of Sec7p, ∼8.0% of Gga2p, and ∼68% of Vps8p signals overlapped with GFP-Vps9p puncta (Fig. 6B, C, and Fig. S4D). In contrast, in the *ent3*Δ *ent5*Δ mutant Vps9p puncta overlapping with Sec7p or Gga2p signals increased to ∼34% or ∼27%, whereas those overlapping with Vps8p decreased to ∼33% (Fig. 6A, D, E, and Fig. S4D). This indicates that transport of Vps9p from the TGN to the Vps21p compartment is suppressed in the *ent3*Δ *ent5*Δ mutant, as shown previously (Nagano et al., 2019). In the *ent3*Δ *ent5*Δ *apl4*Δ mutant GFP-Vps9p puncta overlapping with Vps8-mCH signals were further decreased (∼19%) although those with Sec7-mCherry or Gga2-tdTomato signals were not changed (∼32% or ∼35%), suggesting that AP-1 disruption further suppressed transport of Vps9p from the TGN to the Vps21p compartment (Fig. 6A, D-G and Fig. S4D). Interestingly, GFP-Vps9p puncta overlapping with Vps10-mScarlet-I (Vps10-mSCI), cycling between the TGN and PVC (Casler et al., 2021; Cooper and Stevens, 1996), were significantly decreased in the *ent3*Δ *ent5*Δ *apl4*Δ mutant, compared to the *ent3*Δ *ent5*Δ mutants (∼29% or ∼53%, respectively) (Fig. 6H, I, and Fig. S4E). Thus, in the *ent3*Δ *ent5*Δ *apl4*Δ mutant, GFP-Vps9p seems to be transported from the TGN to some other compartment that is not labeled with these markers (Fig. 6A).

**Figure 6.**
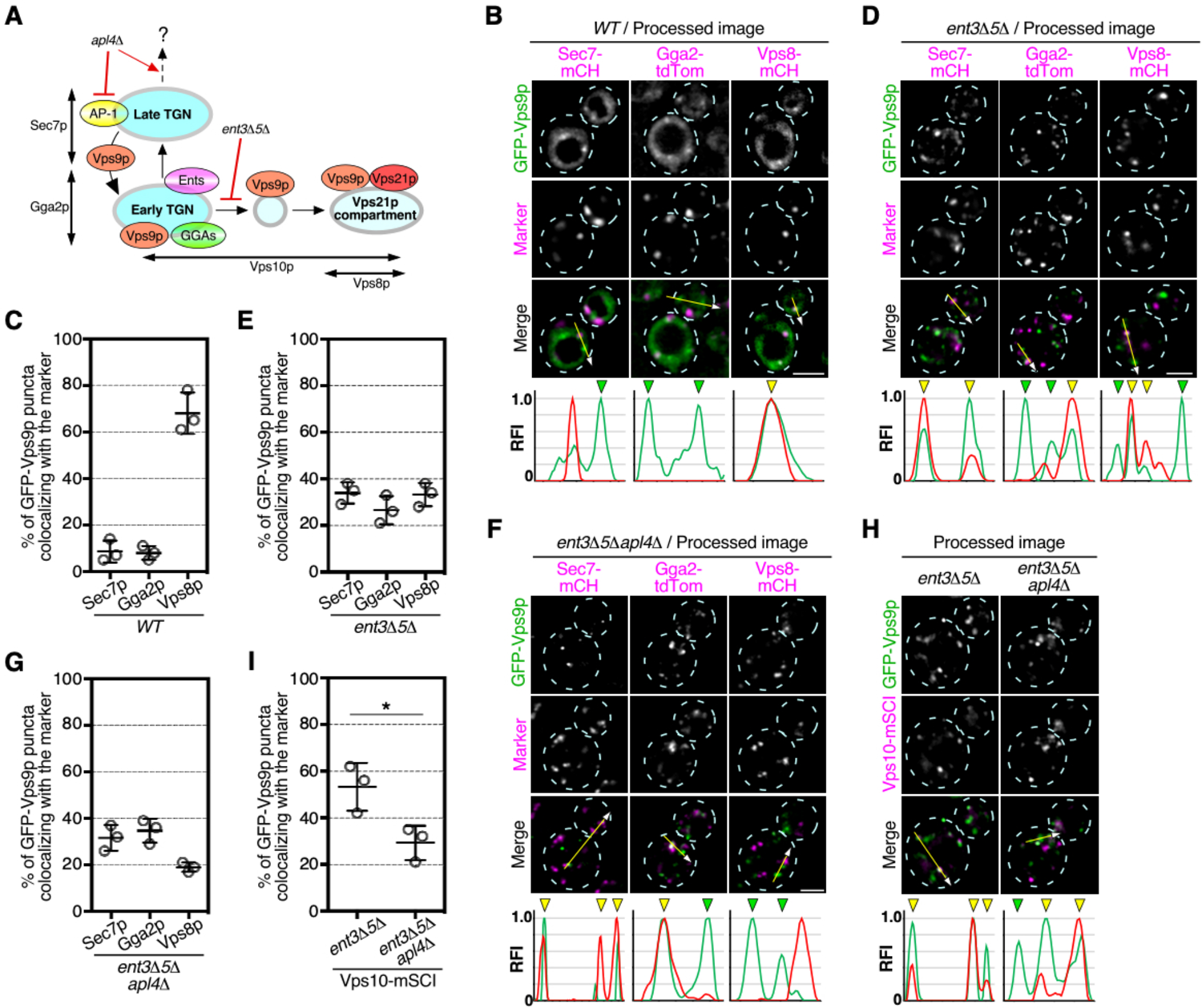
AP-1 complex disruption suppresses the Vps9p loading onto the endosome targeting compartment. (**A**) Schematic illustration of the Vps9p localization at the TGN/endosome and the effect of the disruption of Ent3p/5p and AP-1 complex. Distributions of the maker proteins (Sec7p, Gga2p, Vps10p, and Vps8p) used in Fig. 6 are indicated at the right or lower side. (**B, D, F, H**) Dual color imaging of GFP-Vps9p and red fluorescent protein (RFP)-labeled marker in the *WT* or the indicated mutant cells. Fluorescence images of GFP-Vps9p and the RFP-labeled markers shown in Fig. S4D and E were processed to reduce the cytoplasmic fluorescence background by sequential filtering using Top-hat and bilateral filters bundled in the Image J Fiji software package as shown in Fig. S4A. Representative fluorescence intensity profiles along an arrow are indicated in the right graphs. Yellow or green arrowheads indicate the presence or absence of the overlapping puncta, respectively. (**C, E, G, I**) Quantification of GFP-Vps9p puncta overlapping with the RFP-labeled markers in the cells displayed in (B), (D), (F) or (H). Data show mean ± SEM from three independent experiments in which 100 compartments were scored per each experiment. *p < 0.05, unpaired *t*-test with Welch’s correction. Scale bars: 2.5 μm

## Discussion

The AP-1 complex, GGAs, and epsin-related proteins have been characterized as key factors regulating the transport of clathrin-mediated vesicles from the TGN (Casler and Glick, 2020; Casler et al., 2019; Čopič et al., 2007; Costaguta et al., 2006; Daboussi et al., 2012; Duncan et al., 2003b). In a previous study, we have shown that Ent3p/5p play a crucial role in Rab5 activation (Nagano et al., 2019), but the involvement of AP-1 and GGAs in the process remained unclear. In the present study, we demonstrated that these adaptors play a distinct role in the process of Rab5 activation: Ent3p/5p and AP-1 are required for loading Vps9p on the vesicles that transport Vps9p to the Vps21p compartment, while GGAs, together with Pik1p and Rab11s, have a role in the recruitment of Ent3p/5p and AP-1 to the TGN (Fig. 7A). We have recently reported that the Tlg2p-residing compartment of the TGN, which is distinct from the late TGN where Sec7p resides, is the first destination for endocytic traffic, from which the endocytic cargo is transported to the Vps21p compartment, dependently on GGAs (Toshima et al., 2022). Gga2p is localized predominantly at the Tlg2p-residing compartment (Toshima et al., 2022), and thus Vps9p is probably recruited to and transported from the compartment where GGAs are enriched. In support of this idea, Vps21p-residing vesicles are observed to often make contact with the Tlg2p sub-compartment (Toshima et al., 2022). In the *ent3*Δ *ent5*Δ mutant, efficient loading of Vps9p onto GGA vesicles is likely to be partially impaired (Fig. 7B).

**Figure 7.**
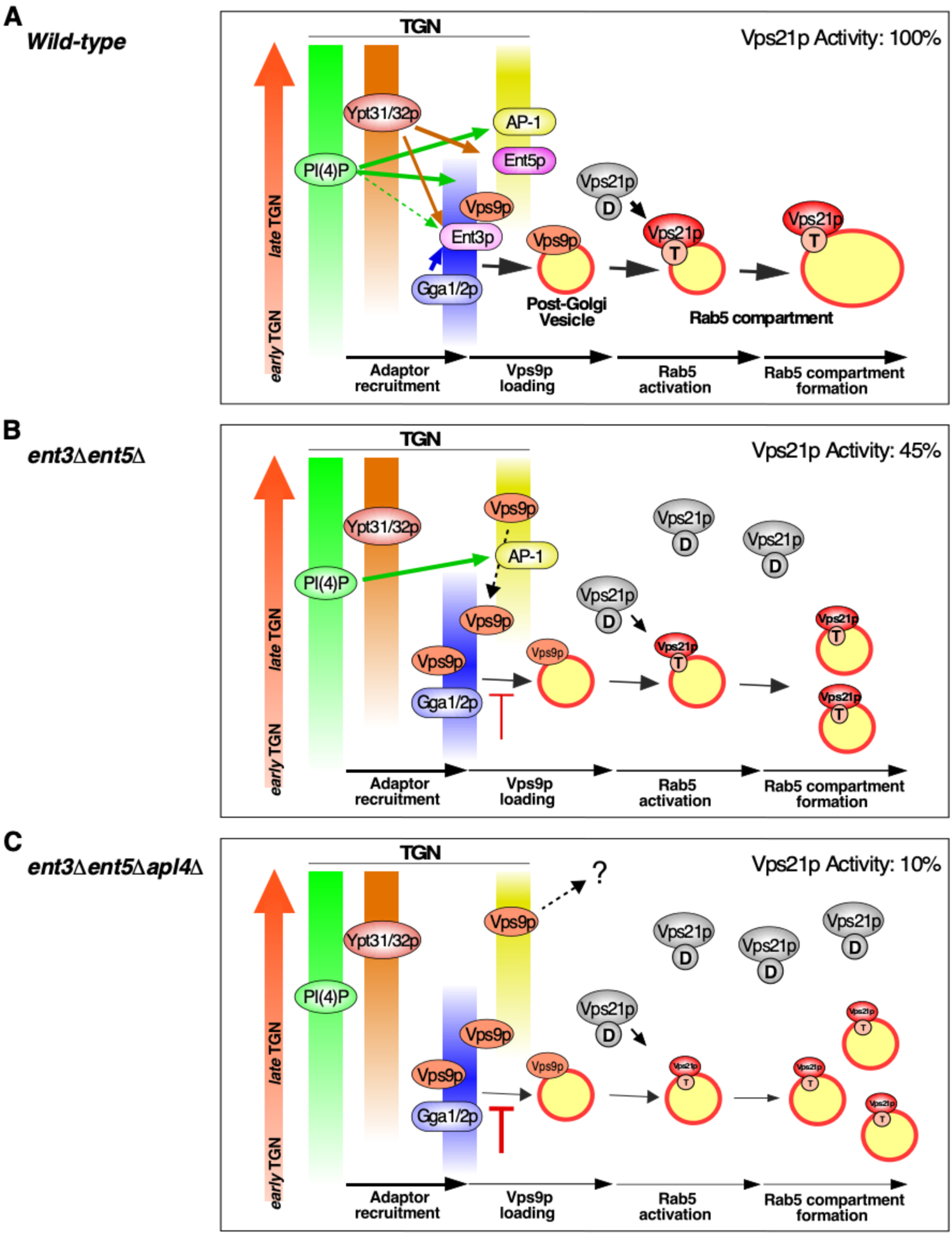
The functional relationships of clathrin adaptors and its upstream regulators in the process of Vps21p activation. (**A**) Overall view of the process of Vps21p activation via GGA and AP-1-enriched regions. Temporal dynamics of PI(4)P, Ypt31p/32p, and the clathrin adaptors are drawn as thick colored lines. GGA- or AP-1-enriched regions are formed sequentially at the late TGN. PI(4)P produced by Pik1p is required for the recruitment of AP-1 and Ent5p, and Ypt31p/32p recruit Ent3p and Ent5p to the TGN. Gga1p/2p also recruits Ent3p to the GGA-enriched region. Efficient recruitment of the clathrin adaptors by these upstream factors facilitates the loading of Vps9p onto the transport vesicles, then Vps9p recruits and activate Vps21p at the vesicles. (**B, C**) Defective formation of Vps9p vesicle in indicated mutant cells. The level of active Vps21p in the mutant relative to that in the wild-type cell was indicated at the upper-right in the panel. See details in the text.

Although deletion of *APL4* had little effect on Vps21p-residing compartment formation, the combination with *ENT3*/*5* deletion resulted in an additive defect. Interestingly, *ENT3*/*5* deletion shifted the localization of Vps9p from endosomes to the TGN, but *APL4* deletion in the *ent3*Δ/*ent5*Δ mutant did not cause an increase in Vps9p localization at the TGN although it decreased localization at endosomes (Fig. 6). Casler et al. have reported that AP-1 cooperates with Ent5p to mediate intra-Golgi recycling of TGN proteins (Casler et al., 2021). In accordance with this model, one possible explanation for the role of AP-1 in Rab5 activation is that AP-1 functions to maintain Vps9p at the TGN, thereby increasing the efficiency of its loading onto transport vesicles. Since in the *ent3*Δ *ent5*Δ mutant the loading of Vps9p onto transport vesicles in the GGAs-enriched region is impaired, the role of AP-1 that recycles Vps9p back to the early TGN might become apparent. Thus, loss of AP-1 in the *ent3*Δ/*ent5*Δ mutant would possibly further reduce the efficiency of Vps9p transport to the Vps21p compartment (Fig.7C). Ent5p, together with AP-1, plays a crucial role in the recycling of TGN-resident proteins, but its role at the GGA-enriched region has remained ambiguous (Casler et al., 2021; Čopič et al., 2007; Costaguta et al., 2006; Daboussi et al., 2012; Hung and Duncan, 2016). We have shown here that Ent5p is able to complement the function of Ent3p in Rab5 activation. This suggests that Ent5p has dual roles at the TGN: loading of Vps9p onto transport vesicles at the GGA-enriched early TGN, and recycling at the AP-1-enriched late TGN.

PI(4)P produced by Pik1p is known to be an important regulator in the recruitment of AP-1, Ent3p/5p, and GGA protein, but the dependence of these adaptors on PI(4)P seems to differ (Daboussi et al., 2012; Duncan and Payne, 2003; Hirst et al., 2003; Wang et al., 2007). Interestingly, our analysis showed that the *gga1*Δ *gga2*Δ mutant had a more severe effect on the localization of Ent3p at the TGN than the *pik1-1* mutant. Thus, although Pik1p, Ypt31p/32p, and GGAs seem to have overlapped roles in the recruitment of Ent3p/5p and AP-1, it is still unclear how these functions are coordinated. Since the function of Arf1p is reportedly related to these proteins, Arf1p is one of the most promising candidates for the coordinator role. Fromme’s group has reported that Ypt31p promotes the activation of Arf1p through its GEF, and that the activated Arf1p recruits the Rab11-GEF TRAPPII complex to promote further activation of Ypt31p/32p (McDonold and Fromme, 2014; Richardson et al., 2012). Similar Arf1p-mediated positive feedback is observed between Pik1p and GGAs (Daboussi et al., 2012). After being recruited to the TGN, Arf1p recruits GGAs, which further functions in Pik1p recruitment to promote PI(4)P production at the TGN. A recent study has demonstrated that the transition stages of TGN are categorized at least tow stages; the early TGN stage which receives retrograde traffic from the endosome, and late TGN stage where transport carriers, such as GGA and AP-1 vesicles, are produced (Tojima et al., 2019). Thus, loading of Vps9p to these transport carriers is presumed to occur at the late TGN stage. Our previous observation showed that treatment of Brefeldin A, which perturbs Golgi maturation by inhibiting Arf1p activity, causes defect in Vps21p-mediated endosome formation (Nagano et al., 2019). These observations suggest that Arf1p is required for both of the TGN progression and Vps21-mediated endosome formation, and regulates the endosome generation coupled with the TGN progression.

In mammalian cell, as well as yeast, interaction of the GGAs and Rabex-5-Rabaptin-5 complex, which is the mammalian orthologue of yeast Vps9p, is reported (Mattera et al., 2003). This study demonstrated that the interaction between GGAs and Rabaptin-5 decreases binding of clathrin to the GGAs, and that overexpression of Rabaptin-5 shifts the localization of GGAs and its associated cargo from the TGN to early endosome (Mattera et al., 2003). These observations suggest a mechanism that the Rab5-GEF, Rabex-5, is recruited to the TGN by GGAs and promotes cargo transport from the TGN to the early endosome via GGA vesicles. Human Rab5 and Rabex-5 have been reported to be upregulated and promote the metastasis ability in several types of malignant tumor cells (Jian et al., 2020; Wang et al., 2014). Intriguingly, mammalian RAB11, ARF1, and PI4KIIIβ (homologues for Arf1p, Rab11 and Pik1p) are also upregulated in these tumor cells (Ferro et al., 2021; Gu et al., 2017; Morrow et al., 2014). These suggests that the TGN-mediated Rab5 activation may be implicated in the tumor invasion and metastasis (Mendoza et al., 2013; Mendoza et al., 2014).

## Materials and methods

### Yeast strains and plasmids

The yeast strains used in this study are listed in Table S1. All strains were grown in standard rich medium (YPD) or synthetic medium (SM) supplemented with 2% glucose and appropriate amino acids. The NH_2_- or COOH-terminal fluorescent protein tagging of proteins was performed as described previously. The plasmids used in this study are listed in Table S2. Fluorescent protein tagging yeast expression constructs were created using pBluescript II (pBS II) vector backbone as previously reported (Nagano et al., 2019). The PCR template plasmids to amplify a gene knockout cassette including *loxP* sequences were created as follows: The PCR fragment of *loxP*-*KlLEU2*-*loxP* or *loxP*-*KlURA3*-*loxP* sequence was amplified from pUG73 (Euroscarf) or pUG72 (Euroscarf), respectively. The fragments were cloned into the pBS II at the EcoRV site. The Cre recombinase yeast expression vector was created as follows: The PCR fragment of the *cre* open reading frame was amplified from pSH67 (Euroscarf). The fragment was cloned into the pGK411 vector (Ishii et al., 2008) (National Bio-Resource Project, Japan) at the SalI and BglII site.

Hexa-Histidine (6×His)-tagged bacterial expression constructs were created using pColdI (Takara) vector backbone. For the assay of active Vps21p quantification, we created the following plasmids: pColdI-GFP nanobody (Katoh et al., 2015; Katoh et al., 2016) (GNB)-Nano luciferase (Boute et al., 2016) (Nluc) for 6×His-GNB-Nluc bacterial expression and pColdI-human Rabenosyn-5 N-terminal (1-40 a.a.) fragment (Qi et al., 2015) (RbNT)-Nano-luciferase (Nluc) for 6×His-RbNT-Nluc. To generate pColdI-GNB-Nluc or pColdI-RbBT-Nluc, a PCR fragment of the Nluc coding sequence amplified from pNL1.1 (Promega) was inserted into the XhoI and EcoRI sites of pColdI to generate pColdI-Nluc. Then, the GNB PCR fragment amplified from pGEX6P1-GNB or RbNT PCR fragment was inserted into the EcoRI and HindIII sites of pColdI-Nluc to generate pColdI-GNB-Nluc or pColdI-hRbNT-Nluc, respectively.

### Fluorescence microscopy

Fluorescence microscopy was performed using an Olympus IX83 microscope equipped with a x100/NA 1.40 (Olympus) objective and an Orca-R2 cooled CCD camera (Hamamatsu), using Metamorph software (Universal Imaging). Simultaneous imaging of red and green fluorescence was performed using an Olympus IX81 microscope, described above, and an image splitter (Dual-View; Optical Insights) that divided the red and green components of the images with a 565-nm dichroic mirror and passed the red component through a 630/50 nm filter and the green component through a 530/30 nm filter. Dual color time lapse imaging of red and green fluorescence was performed using an Olympus IX83 microscope equipped with a high-speed filter changer (Lambda 10-3; Shutter Instruments) that can change filter sets within 40 ms. Images for analysis of colocalization were acquired using simultaneous imaging (64.5 nm pixel size), described above.

### Image processing and analysis

Colocalization between GFP- and mCherry/tdTomato-tagged proteins was quantified by creating masks from GFP and mCherry/tdTomato channels after sequential filtering using the top-hat (Legland et al., 2016) and bilateral (Chaudhury et al., 2011) filters bundled in the Image J FIJI software package (Schindelin et al., 2012). The accuracy of each mask was checked as illustrated in Fig S3. Intensity profiles of GFP-tagged protein and mCherry/tdTomato-tagged protein were generated across the center of fluorescence signals used for the assessment. Colocalization was defined as occurring when the distance between the two peaks of GFP and mCherry/tdTomato intensities was less than 129 nm (2 pixels).

### Recombinant protein purification

The bacterial expression vectors were transformed into the KRX strain (Promega). GST-tagged protein expression was induced by 0.1% rhamnose. The cells were lysed in GST-buffer (20 mM Tris-HCl {pH 8.0}, 100 mM NaCl, 1 mM EDTA, 1 mM DTT, 0.5% NP-40). The protein was purified from the lysate using Glutathione-sepharose 4B (GE Healthcare). His-tagged protein expression was induced by shifting to 15°C and adding 0.1 mM IPTG. The cells were lysed in NP-40 lysis buffer (20 mM Tris-HCl {pH 8.0}, 300 mM NaCl, 20 mM Imidazole, 0.5% NP-40). The protein was purified from the lysate using Ni-NTA agarose (FUJIFILM Wako). Protein purity was checked by SDS-PAGE followed by CBB stain.

### Quantification of active Vps21p level

The method for quantifying active Vps21p levels is illustrated in Fig. 2A. Cells expressing GFP-ALFA-Vps21p were grown in 200 ml YPD to OD^600nm^ of 1.0. The cells were harvested by centrifugation, washed with water, and resuspended in lysis buffer (20 mM Tri-HCl {pH 8.0}, 100 mM NaCl, 10 mM MgCl_2_, protease inhibitor cocktail, 5 mM 1,10-phenanthroline). Glass beads were added to an equal volume and cells were disrupted by Disruptor-Genie (Scientific industry) in the cold room. 600 μg of cleared lysates were incubated with 45∼60 μg of GST-ANB bound to Gluthatione-sepharose 4B for 1 hr in the cold room. The sepharose was washed three times with 5mL wash buffer (20 mM Tri-HCl {pH 8.0}, 500 mM NaCl, 10 mM MgCl_2_, 0.1% NP-40, protease inhibitor cocktail, 5 mM 1,10-phenanthroline), then divided into two aliquot fractions. Next, 10 pmole of 6×His-GNB-Nluc or 6×His-RbNT-Nluc was added into either fraction, then incubated for 1 hr in the cold room. After washing the sepharose six times with 5mL wash buffer, the bound GFP-ALFA-Vps21p was eluted with 200 μg of ALFA peptide. Luciferase activity of the eluted fractions was quantified using the Nano-Glo Luciferase assay system (Promega).

### Western blot assay

Immunoblot analysis was performed as described previously (Toshima et al., 2005). The rabbit polyclonal antibody to GFP (GeneTex, GTX113617) was used at a dilution of 1:10,000 and the HRP-linked donkey F(ab’)_2_ fragment to rabbit IgG (GE Healthcare, NA9340) was used as the secondary antibody at a 1:10,000 dilution. Both mouse monoclonal antibodies to GST (Cell signaling, 26H1) and GAPDH (GeneTex, GTX627408) were used at a dilution of 1:10,000 and the HRP-linked sheep F(ab’)_2_ fragment to mouse IgG (GE Healthcare, NA9310) was used as the secondary antibody at a 1:10,000 dilution. Immunoreactive proteins bands were visualized using the WesternLightning Plus ECL (PerkinElmer).

### Statistics

Statistical analysis was performed with GraphPad Prism 7 software, and the data are shown as the mean ± S.D. or the mean ± S.E.M. as shown in figure legends. Statistical significance was determined using unpaired *t*-test with Welch’s correction or one-way ANOVA with post-hoc Turkey’s test.

### Data availability

The authors declare that all data supporting the findings of this study are available within the article and its supplementary information files.

## Acknowledgements

We thank Dr. Yohei Katoh and Prof. Kazuhisa Nakayama (Kyoto University) for kindly providing the pGEX6P1-GFP nanobody and pGEX6P1-GST-ALFA nanobody vectors. This work was supported by JSPS KAKENHI GRANT #21K06157 to M.N., JSPS KAKENHI GRANT #18K062291, and the Takeda Science Foundation to J.Y.T., as well as JSPS KAKENHI GRANT #19K065710, the Takeda Science Foundation, Life Science Foundation of JAPAN to J.T.

## Author contributions

M. Nagano designed and performed most experiments, analyzed data, and wrote the manuscript. K. Aoshima performed experiments and analysis of *pik1* mutant. H. Shimamura performed experiments and analysis of GFP-Vps21p localization. D.E. Siekhaus reviewed and edited the manuscript, and provided critical input. J.Y. Toshima and J. Toshima designed and supervised the study and wrote and edited the manuscript.

## Supplementary Figure legends

**Figure S1.**
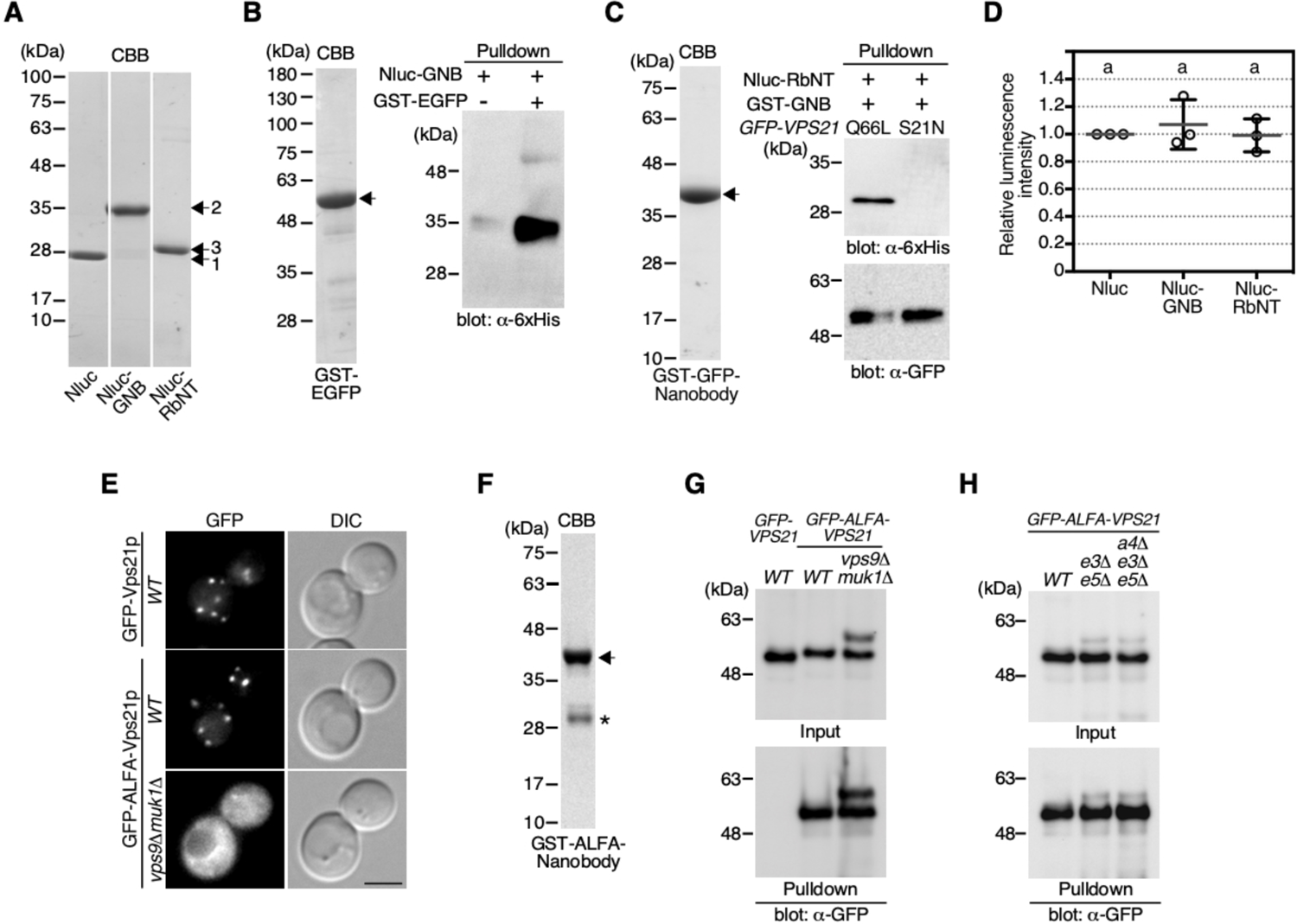
Purification and activity of Nluc-fused probes, and expression of GFP-ALFA-Vps21p. (**A**) Purification of Nluc-fused proteins. Recombinant 6×His-Nano luciferase (Nluc, 1), 6×His-Nluc-GFP nanobody (Nluc-GNB, 2), or 6×His-Nluc-Rabenosyn5 N-terminal fragment (Nluc-RbNT, 3) was expressed in KRX *E. coli* strain, as described in “Material and Methods”. The purified proteins were visualized on SDS-PAGE by Coomassie brilliant blue (CBB) staining. (**B**) Confirmation of the binding ability of Nluc-GNB to GFP. Purified GST-EGFP was visualized by CBB staining (left panel). Nluc-GNB at a final concentration of 200 nM was incubated with Gluthatione sepharose-4B in the presence or absence of the purified GST-EGFP. The bound fractions were subjected to immunoblotting analysis using an anti-6×His antibody (right panel). (**C**) Confirmation of the binding ability of Nluc-RbNT to active Vps21p. Purified GST-GFP nanobody (GST-GNB) was visualized by CBB staining (left panel). The extract of yeast cells expressing the GFP-tagged active Vps21p mutant (Q66L) or the inactive mutant (S21N) were incubated with the GST-GNB-bound Gluthatione sepharose-4B. The sepharose was washed, and then incubated with Nluc-RbNT at a final concentration of 200 nM. The bound fractions were subjected to immunoblotting analysis using an anti-6×His antibody or anti-GFP antibody (right panel). (**D**) Relative luminescence intensity of the probes shown in (A). Graph shows the activity of each 100 pM probes. Data show mean ± SEM from three independent experiments. Different letters indicate significant difference at p < 0.05, one-way ANOVA with Tukey’s post-hoc test. (**E**) Expression of GFP-Vps21p or GFP-ALFA-Vps21p in the cells. Scale bar, 2.5 μm. (**F**) Purified GST-tagged ALFA nanobody was visualized by CBB staining (**G, H**) Immunoblots showing the amount of GFP-Vps21p or GFP-ALFA-Vps21p in the cell extract (Input) or the pulldown fraction (Pulldown).

**Figure S2.**
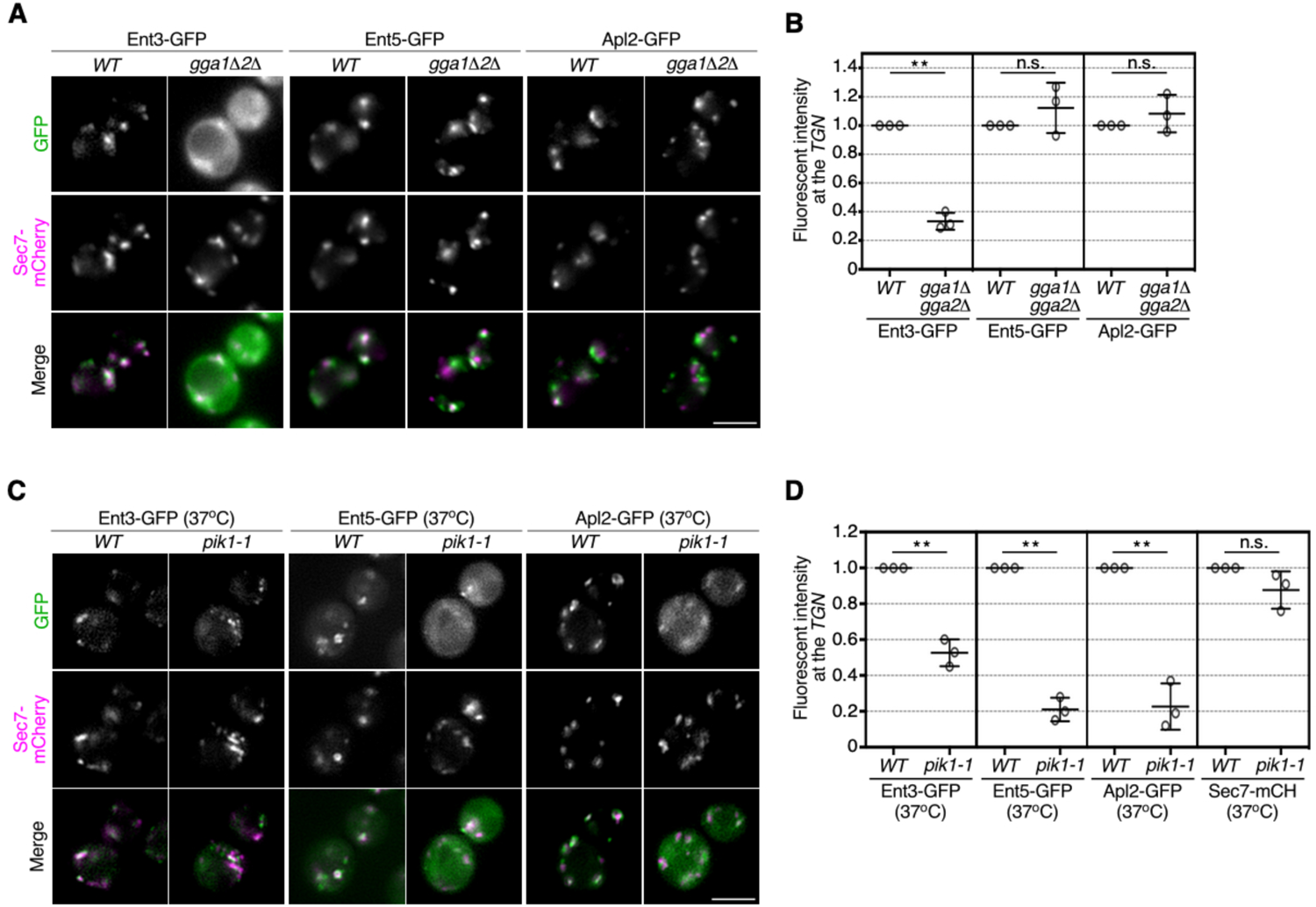
The effect of GGA and Pik1p disruption on the clathrin adaptor localization. (**A**) Localization of GFP-tagged Ent3p (Ent3-GFP), Ent5p (Ent5-GFP), or Apl2p (Apl2-GFP) in the *gga1*Δ*2*Δ cells. Sec7-mCH was expressed as a marker for the TGN. (**B**) Relative fluorescence intensity of Ent3-GFP, Ent5-GFP, and Apl2-GFP at the TGN in the *gga1*Δ*2*Δ cells. (**C**) Localization of Ent3-GFP, Ent5-GFP, and Apl2-GFP in *pik1-1* temperature-sensitive mutant cells. Sec7-mCH was expressed as a control to evaluate the effect of Pik1p dysfunction on Golgi/TGN function. Pik1p function was diminished by incubating cells at 37 °C for 2 h. (**D**) Quantification of the fluorescence intensity of Ent3-GFP, Ent5-GFP, and Apl2-GFP at the TGN (as labeled by Sec7-mCH) in *pik1-1* cells. Intensity of Sec7-mCH was used as a control. Data show mean ± SEM from three independent experiments in which 50 puncta were scored per each experiment. **p < 0.01, n.s., not significant, unpaired *t*-test with Welch’s correction.

**Figure S3.**
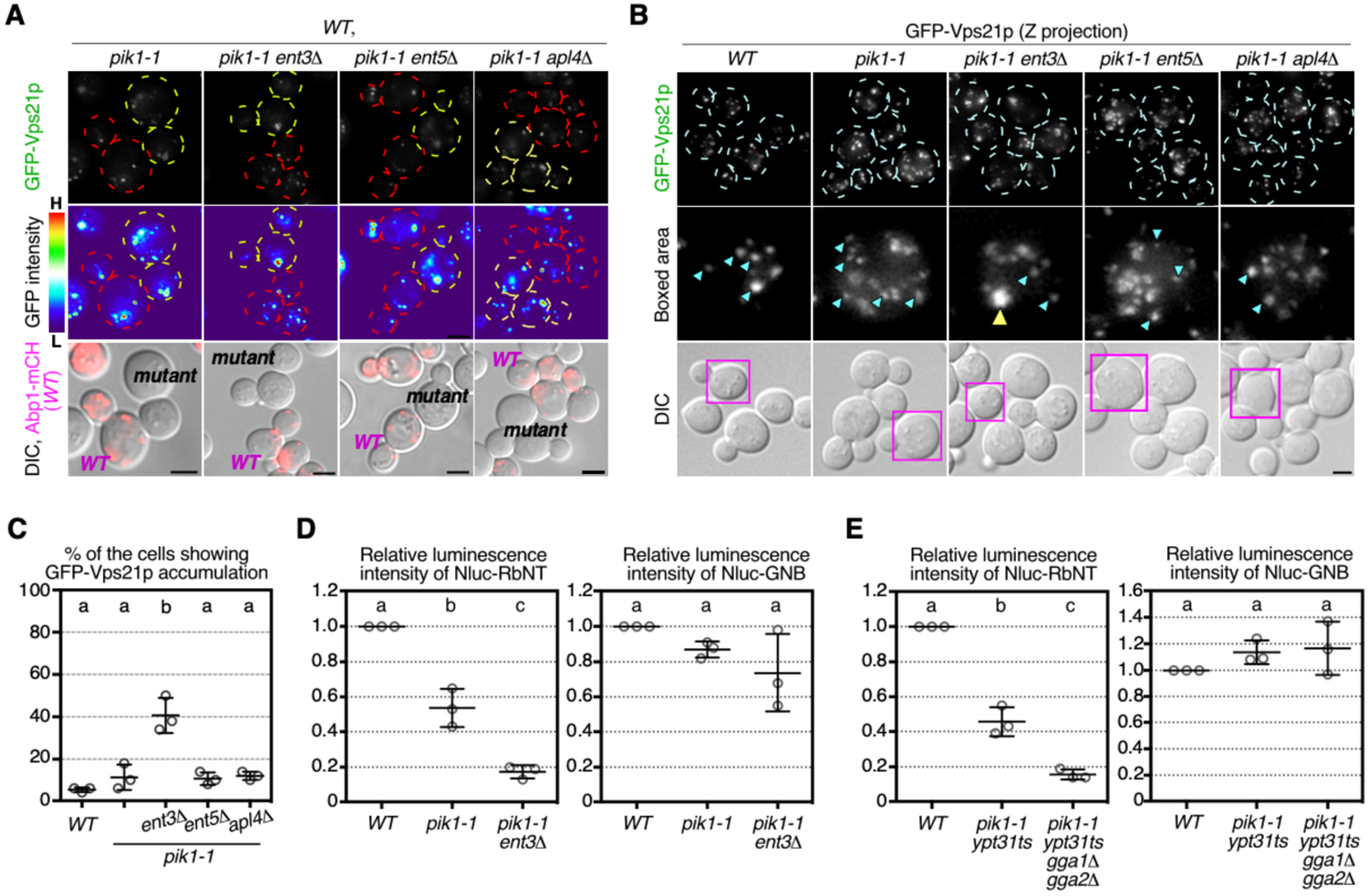
Pik1p is involved in the Vps21p activation. (**A**) Localization of GFP-Vps21p in the *pik1-1* or the clathrin adaptor-defective *pik1-1* mutants. Pik1p function was diminished by incubating cells at 37 °C for 2 h. Fluorescence images or heat maps showing GFP levels are shown in the panels labeled GFP-Vps21p or GFP intensity, respectively. The *WT* cells are labeled by the expression of Abp1-mCH (magenta) overlaid with DIC images as shown in the lower row. Red or yellow outline indicate the wild-type or mutant cells, respectively. (**B**) Maximum intensity projections of Z stacks of wild-type (*WT*) and mutant cells expressing GFP-Vps21p. The Z series was acquired through the entire cell at 0.4 μm intervals. Cyan and yellow arrowheads indicate the examples of GFP-Vps21p-residing endosome-like compartments and the GFP-Vps21p-accumulated regions, respectively. Higher magnification view of the boxed area is displayed in the middle panel. **(C)** The percentages of the cells showing the GFP-Vps21p accumulation displayed in (B). (**D, E**) Quantification of the binding amount of Nluc-fused probes in the total or active Vps21p pulldown fraction. GFP-Vps21p or GFP-ALFA-Vps21p was expressed in the indicated cells. According to the protocol illustrated in Fig. 2A, Nluc activity in the Nluc-RbNT-bound active Vps21p fraction (left graph) or the Nluc-GNB-bound total Vps21p fraction (right graph) were quantified. Graphs represent the active Vps21p level in the mutant cells relative to that in *WT* cells. Data show mean ± SEM from three independent experiments in which 50 cells (C) were scored per each experiment, or mean ± SEM from three independent experiments (D, E). Different letters indicate significant difference at p < 0.05, one-way ANOVA with Tukey’s post-hoc test. Scale bar, 2.5 μm.

**Figure S4.**
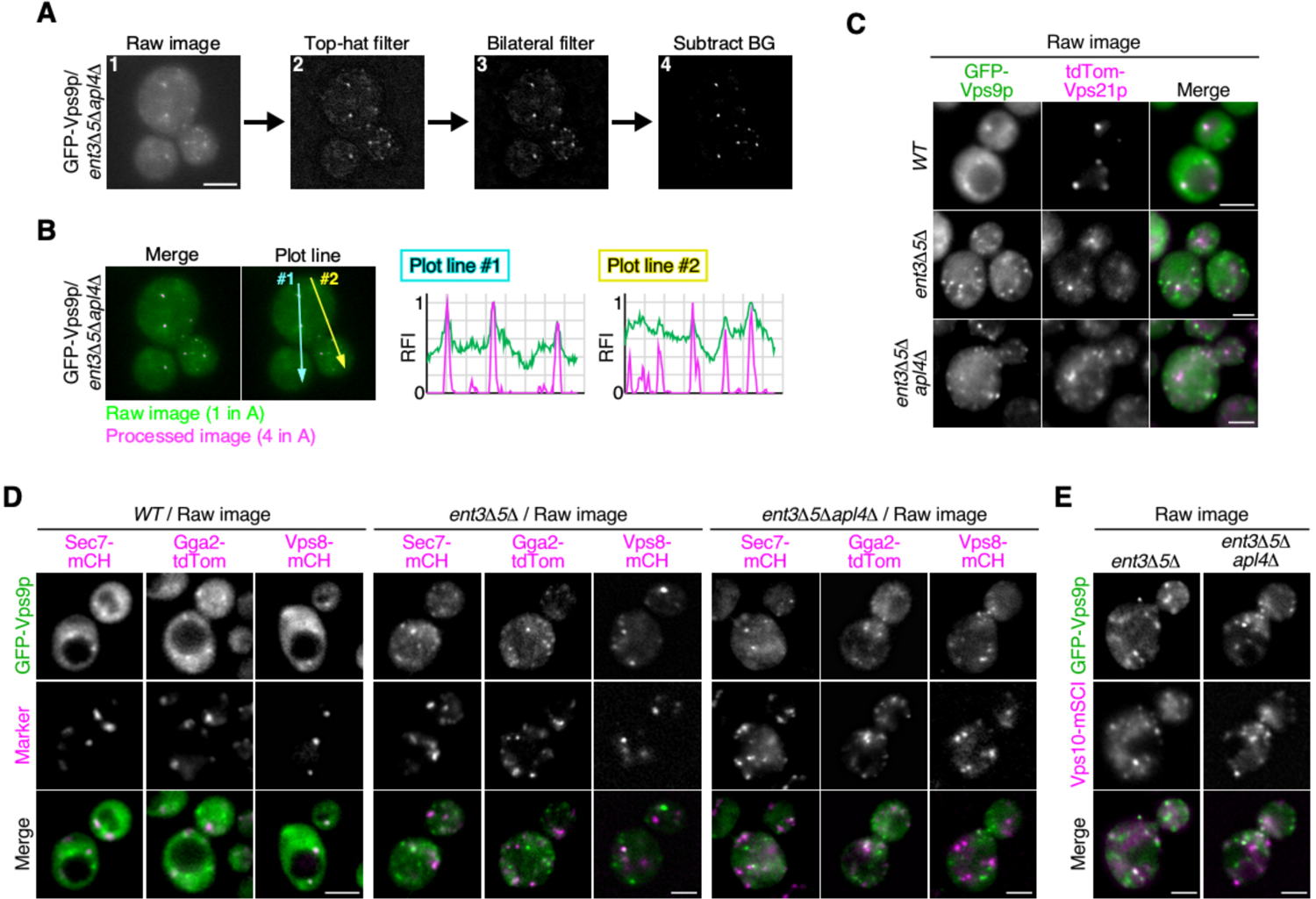
Localization of GFP-Vps9p between TGN and endosomal compartment in the *ent3*Δ*5*Δ and *ent3*Δ*5*Δ *apl4*Δ cells. **(A, B)** Manifestation of GFP-Vps9 fluorescence signals in cell. Steps for the image processing to reduce the cytoplasmic fluorescence background by the sequential Top-hat and bilateral filters (ImageJ FIJI) are indicated in (A). The raw image of cell expressing Vps9p (panel 1) was processed sequentially by the top-hat filter (panel 2), the bilateral filter (panel 3), and smoothing process (panel 4), using software bundled in Image J FIJI. Overlay of raw image and background processed image, and comparison of their fluorescence intensity peaks are indicated in (B). The raw image (panel 1 in A) was overlaid with the final image (panel 4 in A). Representative fluorescence intensity profiles along a line are indicated in the right side. (**C**) Localization of GFP-Vps9p and mCherry-Vps21p in the *ent3*Δ*5*Δ and *ent3*Δ*5*Δ *apl4*Δ cells. The raw images used for image processing in Fig. 5F. Cells expressing GFP-Vps9 and mCherry-Vps21p were grown to log phase at 25°C, and imaged. (D, E) Localization of GFP-Vps9p between TGN and endosomal compartment in the *ent3*Δ*5***Δ** and *ent3*Δ*5*Δ *apl4*Δ cells. The raw images used for image processing in Fig. 6B, D, F are indicated in (D), and that in Fig. 6H are indicated in (E).

---

**Supplementary Table1.**
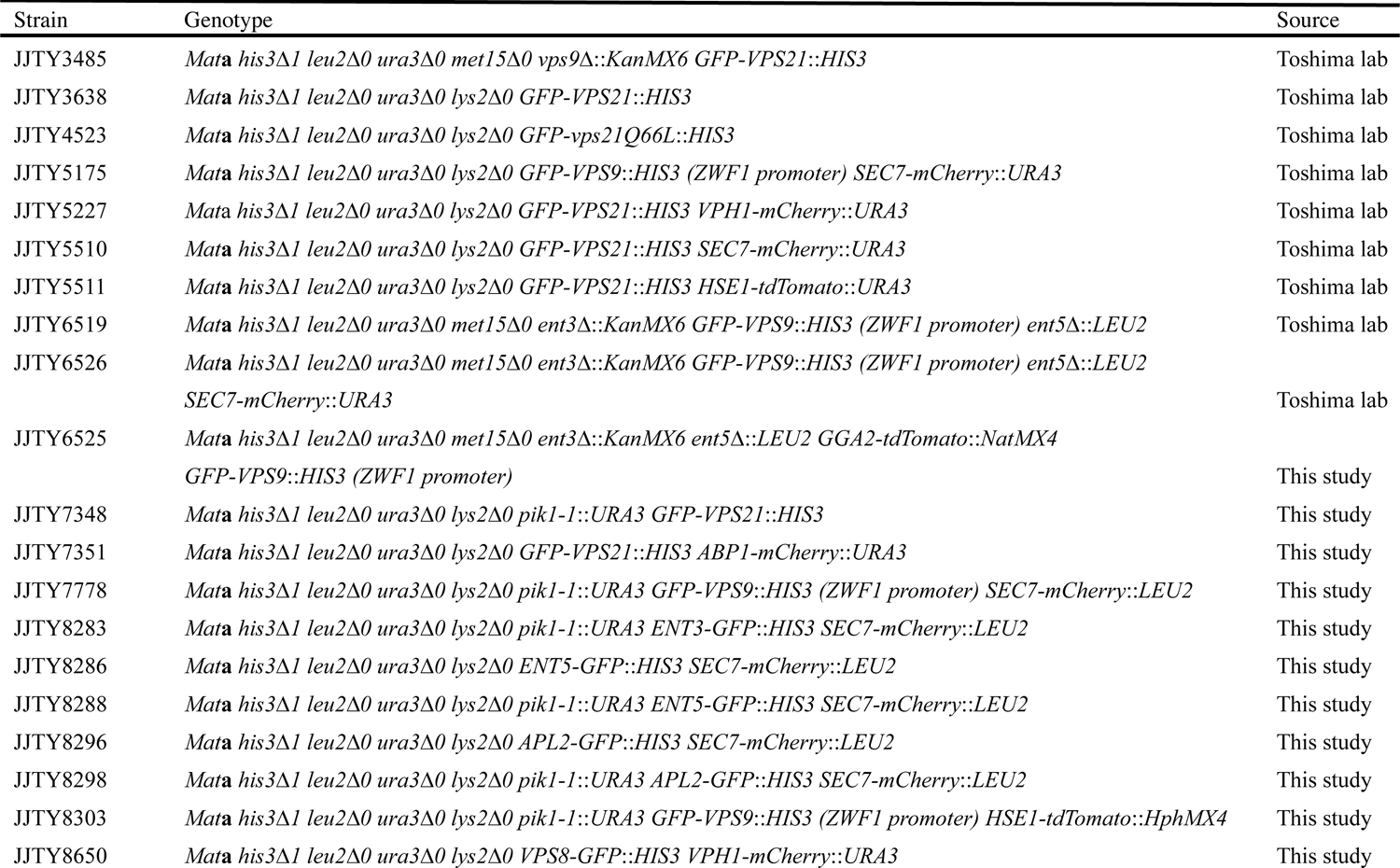
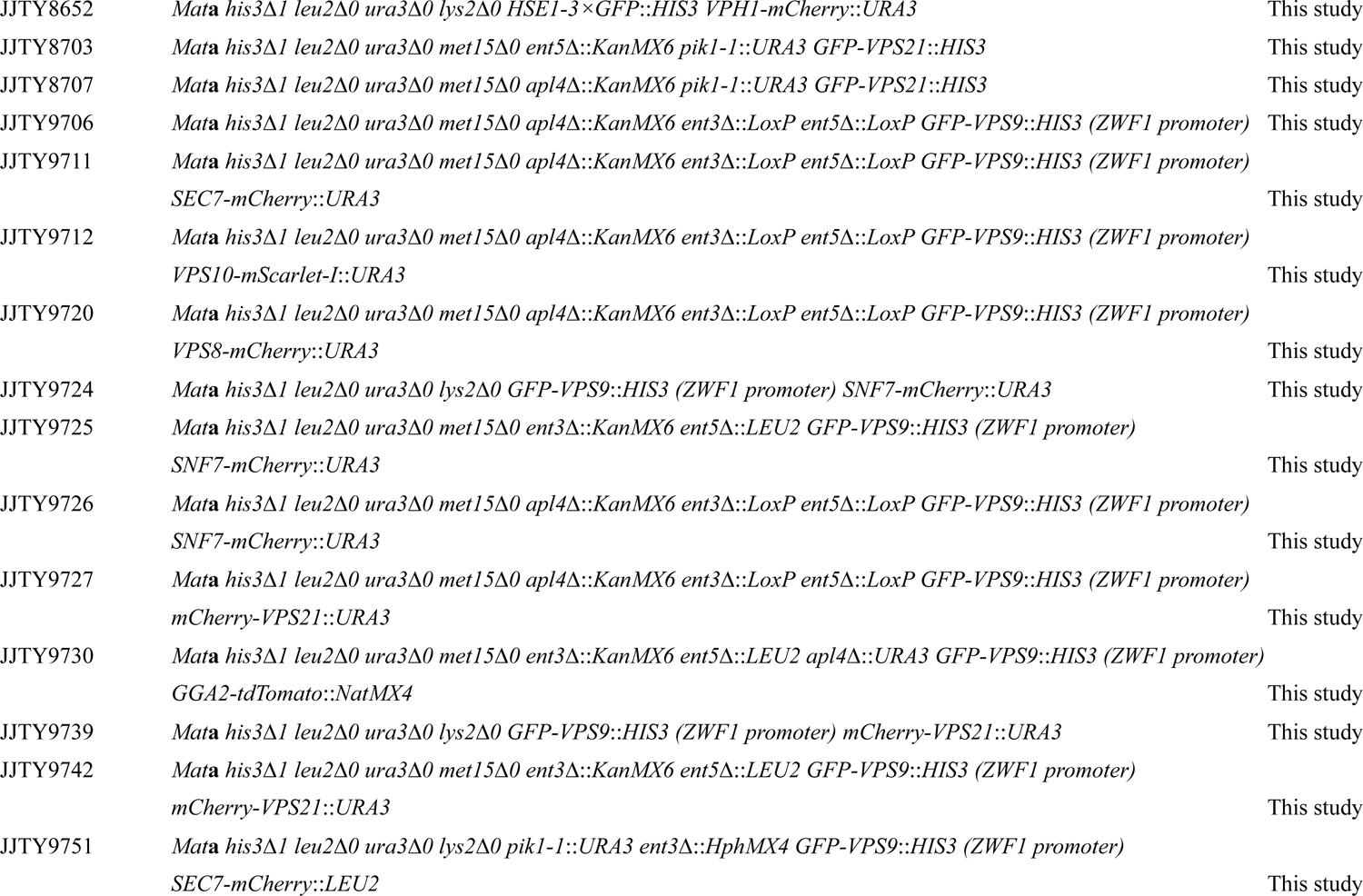
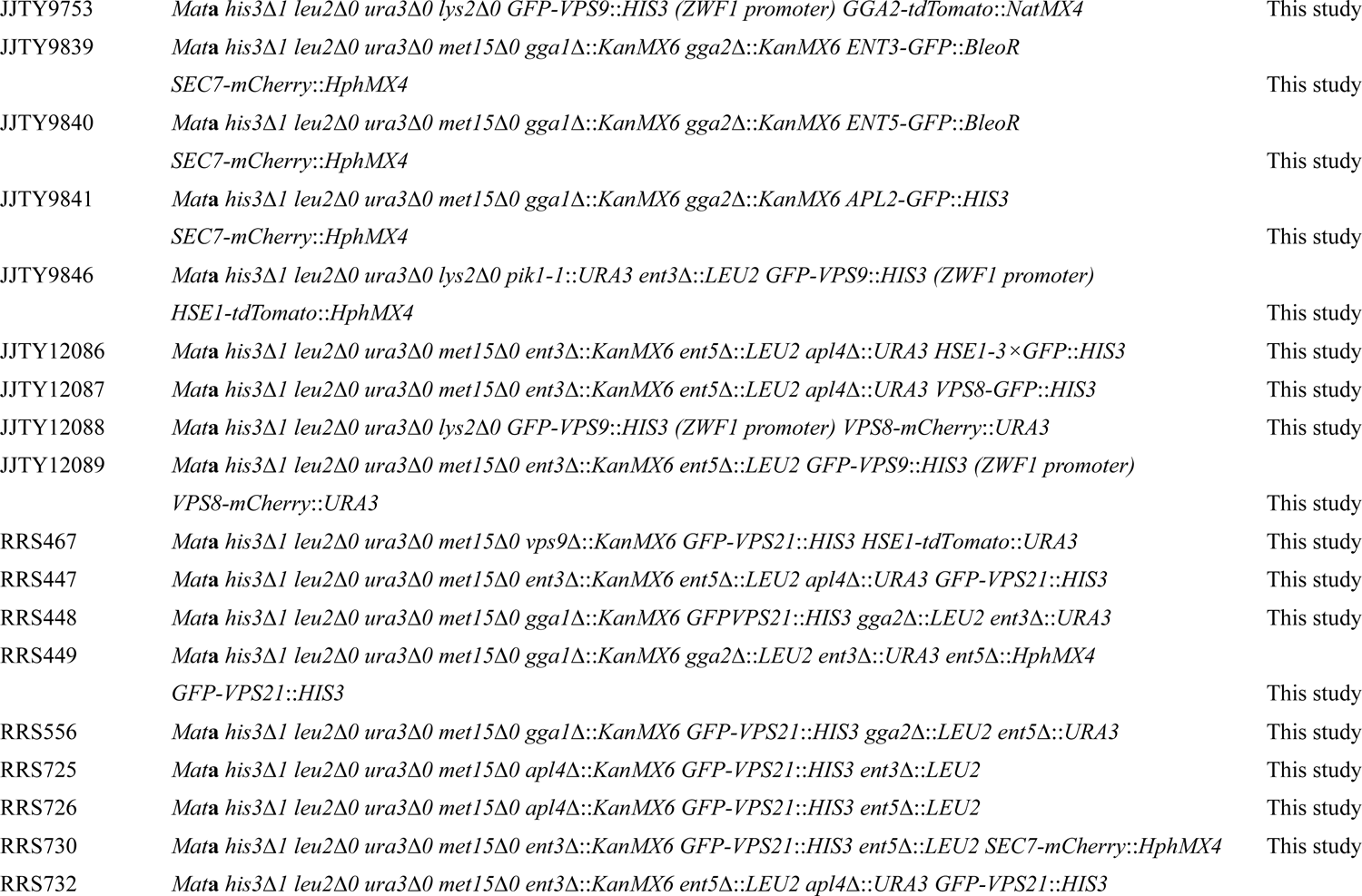
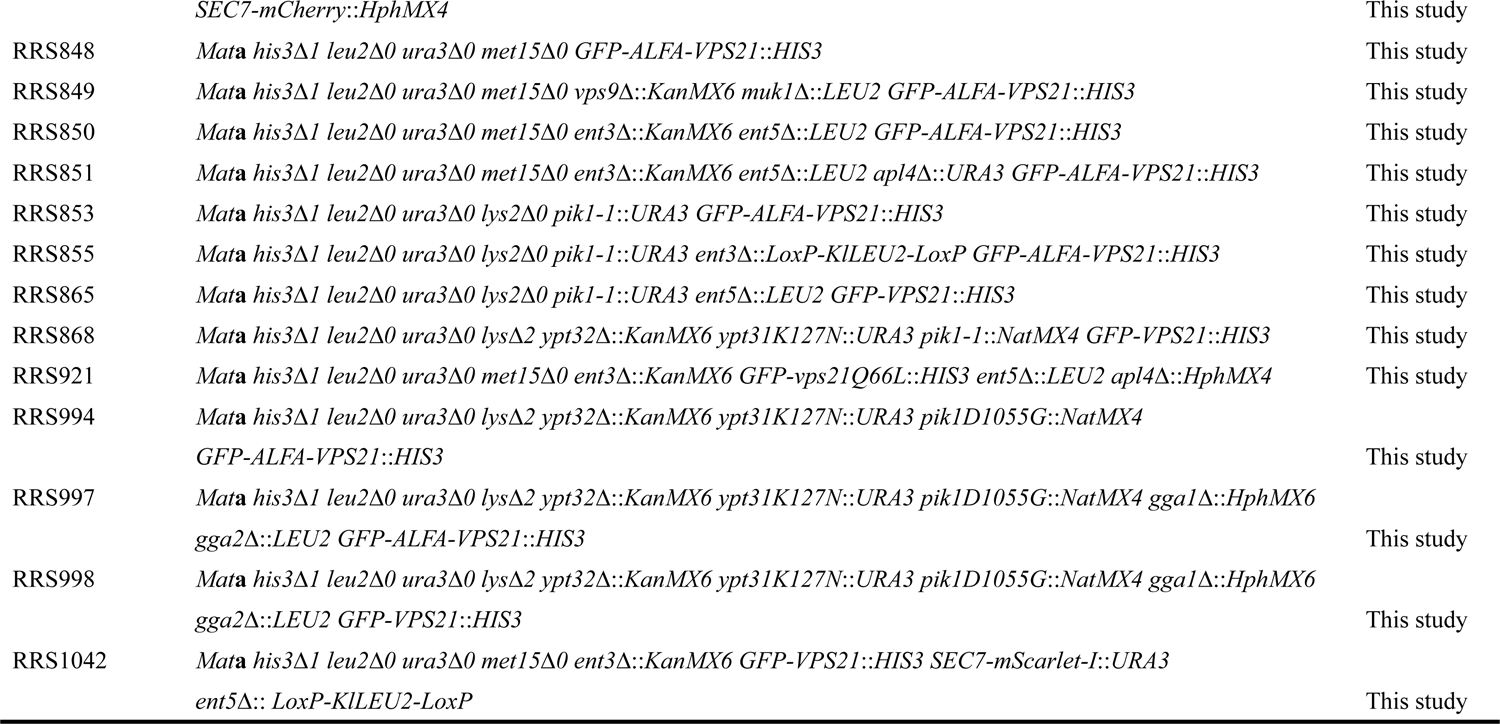
Yeast Strains used in this study

**Supplementary Table2.** Plasmid used in this study

| Name | Purpose | Backbone | Description (for selection) | Source |
| --- | --- | --- | --- | --- |
| For Yeast genetics |  |  |  |  |
| pJT6 | C-terminal GFP tagging | pFA6a | PCR template (DO-HIS) | Toshima lab |
| pJT33 | Gene deletion using *LEU2MX6* | pBS II | PCR template (DO-LEU) | Toshima lab |
| pJT36 | Gene deletion using *URA3MX6* | pBS II | PCR template (DO-URA) | Toshima lab |
| pJT601 | C-terminal mCherry tagging | pFA6a | PCR template (DO-URA) | Toshima lab |
| pJT694 | C-terminal mCherry tagging | pFA6a | PCR template (DO-LEU2) | Toshima lab |
| pJT740 | Hse1p C-terminal 3 × GFP tagging | pBS II | *HSE1-3 × GFP* integration (DO-HIS) | Toshima lab |
| pJT758 | Vps21p N-terminal GFP tagging | pBS II | *GFP-VPS21* integration (DO-HIS) | Toshima lab |
| pJT791 | Vps21Q66L N-terminal GFP tagging | pBS II | *GFP-vps21Q66L* integration (DO-HIS) | Toshima lab |
| pJT792 | Vps21S21N N-terminal GFP tagging | pBS II | *GFP-vps21S21N* integration (DO-HIS) | Toshima lab |
| pJT844 | Vps21p N-terminal mCherry tagging | pBS II | *mCherry-VPS21* integration (DO-URA) | Toshima lab |
| pJT1602 | *ypt31ts* strain production | pBS II | *ypt31K127N* mutagenesis (DO-URA) | Toshima lab |
| pJT1756 | Gene deletion using *HphMX4* | pBS II | PCR template (Hygromycin B) | Toshima lab |
| pJT1878 | Hse1p C-terminal tdTomato tagging | pBS II | *HSE1-tdTomato* integration (DO-URA) | Toshima lab |
| pJT1914 | Vps9p N-terminal GFP tagging | pBS II | *ZWF1 pro-GFP-VPS9* integration (DO-HIS) | Toshima lab |
| pJT2203 | Gene deletion using *NatMX4* | pBS II | PCR template (Nourseothricin) | Toshima lab |
| pJT2205 | Hse1p C-terminal tdTomato tagging | pBS II | *HSE1-tdTomato* integration (Hygromycin B) | Toshima lab |
| pJT2371 | *pik1-1* strain production | pBS II | *pik1D1055G* mutagenesis (DO-URA) | Toshima lab |
| pJT1733 | Gene deletion using *LoxP-KIURA3-LoxP* | pBS II | PCR template (DO-URA) | This study |
| pJT1882 | Gene deletion using *LoxP-KILEU2-LoxP* | pBS II | PCR template (DO-LEU) | This study |
| pJT2188 | Gga2p C-terminal tdTomato tagging | pBS II | *GGA2-tdTomato* integration (Nourseothricin) | This study |
| pJT2823 | C-terminal mScarlet-I tagging | pFA6a | PCR template (DO-URA) | This study |
| pJT2829 | Cre recombinase expression | pGK411 | CEN/ARS vector (DO-MET) | This study |
| pJT2839 | C-terminal GFP tagging | pFA6a | PCR template (Phleomycin) | This study |
| pJT2842 | *pik1-1* strain production | pBS II | *pik1*D1055G mutagenesis (Nourseothricin) | This study |
| pJT2856 | Vps21p N-terminal GFP-ALFA tagging | pBS II | GFP-ALFA-VPS21 integration (DO-HIS) | This study |
| ..... |  |  |  |  |
| For *E. coli* expression |  |  |  |  |
| pJT2749 | GST-GFP nanobody expression | pGEX6P1 | (Ampicillin) | This study |
| pJT2751 | GST-EGFP expression | pDEST15 | (Ampicillin) | This study |
| pJT2831 | 6×His-Nluc-GFP nanobody expression | pColdI | (Ampicillin) | This study |
| pJT2855 | 6×His-Nluc-hRabenosyn 5 (1-40aa) expression | pColdI | (Ampicillin) | This study |
| pJT2899 | GST-ALFA nanobody expression | pGEX6P1 | (Ampicillin) | This study |

